# Interspecies transfer of syntenic *RAMOSA1* orthologs and promoter *cis* sequences impacts maize inflorescence architecture

**DOI:** 10.1101/2022.01.07.475427

**Authors:** Josh Strable, Erica Unger-Wallace, Alejandro Aragón Raygoza, Sarah Briggs, Erik Vollbrecht

## Abstract

Grass inflorescences support floral structures that each bear a single grain, where variation in branch architecture directly impacts yield. The maize RAMOSA1 (*Zm*RA1) transcription factor acts as a key regulator of inflorescence development by imposing branch meristem determinacy. Here, we show *RA1* transcripts accumulate in boundary domains adjacent to spikelet meristems in *Sorghum bicolor* (*Sb*) and *Setaria viridis* (*Sv*) inflorescences similar as in the developing maize tassel and ear. To evaluate functional conservation of syntenic *RA1* orthologs and promoter *cis* sequences in maize, sorghum and setaria, we utilized interspecies gene transfer and assayed genetic complementation in a common inbred background by quantifying recovery of normal branching in highly ramified *ra1-R* mutants. A *ZmRA1* transgene that includes endogenous upstream and downstream flanking sequences recovered normal tassel and ear branching in *ra1-R*. Interspecies expression of two transgene variants of the *SbRA1* locus, modeled as the entire endogenous tandem duplication or just the non-frameshifted downstream copy, complemented *ra1-R* branching defects and induced novel fasciation and branch patterns. The *SvRA1* locus lacks conserved, upstream noncoding *cis* sequences found in maize and sorghum; interspecies expression of an *SvRA1* transgene did not or only partially recovered normal inflorescence forms. Driving expression of the *SvRA1* coding region by the *ZmRA1* upstream region, however, recovered normal inflorescence morphology in *ra1-R*. These data leveraging interspecies gene transfer suggest that *cis*-encoded temporal regulation of *RA1* expression is a key factor in modulating branch meristem determinacy that ultimately impacts grass inflorescence architecture.

## INTRODUCTION

Understanding the genetic basis of morphological diversity between and within species is a key objective in biology (Carroll, 2008). Grass (Poaceae) inflorescences display tremendous intra- and interspecific variation (ref. Kellogg, 2015) and are an effective model for studying genetic mechanisms that underly evolutionary change in morphology. Inflorescence diversity is well-documented in the cereal crops rice (*Oryza* spp.) (Yamaki et al., 2010; Crowell et al., 2016), millet (*Setaria* spp.) (Doust and Kellogg, 2002; Doust et al., 2005; Huang and Feldman, 2017), sorghum (*Sorghum* spp.) (Harlan and de Wet, 1972; Brown et al., 2006; Zhou et al., 2019; Li et al., 2020) and maize (*Zea mays* sspp.) (Upadyayula et al., 2006a; Upadyayula et al., 2006b; Brown et al., 2011; Wu et al., 2016; Xu et al., 2017). As inflorescences in the Poaceae ultimately support reproduction and the floral structures that bear a single grain, variation in inflorescence morphology directly impacts yield in cereal crops and weedy grass species. Despite such agronomical and ecological significance, the genes that underlie diverse inflorescence forms in the grasses have not been fully elucidated, and tests of functional conservation of syntenic orthologous genes are limited.

Mature inflorescence traits are patterned early in development through variation in size, identity, and the timing and duration of maturation schedules of active pluripotent stem cell tissues called meristems. These variations impact the number, arrangement and elaboration of lateral organs that arise from meristems (Doust and Kellogg, 2002; Vollbrecht et al., 2005; Prusinkiewicz et al., 2007; Whipple et al., 2010; Kellogg et al., 2013; Lemmon et al., 2016; Zhu et al., 2018; Leiboff and Hake, 2019). A general framework for ontogeny of grass inflorescences (Kellogg et al., 2013) follows: When internal and external cues signal the reproductive transition, inflorescence development ensues as a vegetative shoot apical meristem, which elaborates leaf primordia at its flanks, converts to a reproductive inflorescence meristem (IM) that elaborates lateral meristems at its flanks. The IM is indeterminate, i.e. capable of producing an unspecified number of lateral primordia, and the lateral meristems can be either relatively indeterminate in which case they may also initiate additional lateral meristems, or relatively determinate (producing a specified number of lateral primordia). Indeterminate grass inflorescence meristems are called branch meristems (BMs) and show diverse indeterminacy across and even within grass species, while all grass inflorescences ultimately produce determinate meristems called spikelet meristems (SMs). Thus, in the general framework IMs initiate BMs, and both IMs and BMs initiate SMs at their flanks. A SM gives rise to two glume (bract) primordia, followed by one or multiple florets which altogether comprise the spikelet, the central unit of a grass inflorescence (Clifford, 1987). SMs in some grass species are more determinate in that they terminate by converting to a floral meristem that is consumed in the production of floral organs, whereas in other species SMs are somewhat indeterminate and produce multiple floral meristems, and therefore multiple florets, before terminating. Diverse morphological complexity among grass inflorescences arises though variation in type, activity and determinacy of IMs, BMs and SMs.

The family Poaceae consists of over 11500 species (Kellogg, 2015) distributed about equally among two major lineages known as the PACMAD and BOP clades. In the PACMAD clade, the largest subfamily Panicoideae has over 3300 species that include global staple cereal crops maize (*Zea mays* ssp *mays*), sorghum (*Sorghum bicolor* [L.] Moench), and foxtail millet (*Setaria italica*) (Kellogg, 2015). Maize and sorghum are among the ∼1200 species in tribe Andropogoneae; *Setaria* is in the tribe Paniceae (Kellogg, 2015). Unlike most of the Panicoideae where spikelets are unpaired, the Andropogoneae are distinguished by producing their spikelets in pairs; specialized, determinate BMs called spikelet pair meristems (SPMs) each produce two SMs. Thus, spikelet pairs (SPs) and long branches (LBs), which commonly coexist in the same inflorescence, are branches that differ by length (short vs. long, respectively) and meristem determinacy at origin (SPMs vs. BMs, respectively). By contrast, within the tribe Paniceae or the ‘bristle clade’ are a few hundred grass species including the foxtail millet progenitor *Setaria viridis* where adjacent meristems differentiate into either single spikelets or sterile branches called bristles (Doust and Kellogg, 2002, Hodge and Doust, 2017). Developmental and morphological studies in *Setaria* lend support to the ontogenetic pairing of a single spikelet with a bristle, but spikelets are not paired (Doust and Kellogg, 2002).

Maize and sorghum are estimated to have diverged from a common ancestor approximately 12 million years ago (MYA) (Swigonová et al., 2014); *Setaria* diverged from maize and sorghum approximately 26-27 MYA (Bennetzen et al., 2012; Zhang et al., 2012). Sorghum and *Setaria* genomes show extensive synteny (Bennetzen et al., 2012; Zhang et al., 2012). Likewise, approximately 60% of annotated genes are syntenically conserved between maize and sorghum, and this gene set accounts for 90% of all genes characterized by forward genetics in maize (Schnable and Freeling, 2011; Schnable, 2015). Syntenic orthologs are more likely to retain consistent patterns of gene regulation and expression across related species (Davidson et al., 2012), and may be more likely to retain ancestral functional roles than non-syntenic gene copies (Dewey, 2011). However, to date, functional conservation between syntenic orthologs in related grass species remains widely untested.

The maize *RAMOSA1* (*ZmRA1*) locus is a key regulator of tassel and ear development and morphology (Vollbrecht et al., 2005). *ZmRA1* was a target of selection during maize domestication (Sigmon and Vollbrecht, 2010), co-localizes with nucleotide polymorphisms for inflorescence branching traits in genome wide association studies of diverse maize breeding lines (Brown et al., 2011; Wu et al., 2016; Xu et al., 2017) and is a candidate quantitative trait locus for tassel branch number in the Mexican highland maize landrace Palomero Toluqueño (Perez-Limón et al., 2021). Strong maize *ra1* mutants were recognized over a century ago as resembling inflorescences of other grasses (Collins, 1917), and more recently, comparisons to the complexly branched sorghum panicle have been drawn at developmental and molecular levels (Vollbrecht et al., 2005; Leiboff and Hake, 2019). Whereas normal inflorescence branching in maize produces only SPs or LBs bearing SPs, mutations in *ZmRA1* relax the determinacy normally imposed on SPMs such that SPs are replaced by LBs bearing several unpaired, single spikelets (“spikelet multimers”), or by LBs bearing a mix of single and/or paired spikelets (Vollbrecht et al., 2005). The graded, multiple orders of inflorescence branching in ra1 mutants reveal a general determinacy function of *ZmRA1* in addition to or that includes a specific role for *ZmRA1* activity in producing the canonical SP. *RA1* encodes a C_2_H_2_ zinc-finger transcription factor with EAR repression motifs (Vollbrecht et al., 2005). Mutations in the maize C_2_H_2_ zinc-finger domain or C-terminal EAR motif result in severe *ra1* mutants that display highly ramified tassels and ears (Vollbrecht et al., 2005; Gallavotti et al., 2010). One mechanism by which RA1 imposes SPM determinacy in maize is through genetic and physical interactions with the orthologous TOPLESS co-repressor encoded by *RA1 ENHANCER LOCUS2* (*REL2*) (Gallavotti et al., 2010). *ZmRA1* transcripts and *Zm*RA1 protein accumulate in a boundary domain between the inflorescence or branch axis and the determinate meristems it regulates (Vollbrecht et al., 2005; Eveland et al., 2014). The non-cell-autonomous nature of *ZmRA1* suggests that it regulates a trafficable signal for meristem determinacy, or its gene product is capable of trafficking to the adjacent meristem (Vollbrecht et al., 2005). Genetic and molecular data support that *RA1* expression in maize impacts branch complexity through regulating SPM determinacy (Vollbrecht et al., 2005). Variation in timing of *RA1* expression, presumably imposed by variation in promoter *cis* sequences, in *Miscanthus* (Vollbrecht et al., 2005), sorghum (Vollbrecht et al., 2005; Leiboff and Hake, 2019) and *S. viridis* (Zhu et al., 2018) correlates with degree of branch activity and distinct inflorescence morphologies. Thus, heterochronic *RA1* expression and regulation of RA1 activity are hypothesized to impact inflorescence branching directly by modulating meristem determinacy. To date, *ra1* mutants have not been reported outside of maize, leaving open the question of *RA1* function in the Panicoideae with respect to evolutionarily and agronomically important characters such as meristem determinacy, branch length and pairing of spikelets.

Here, we report on genetic tests for functional conservation of syntenic orthologous *RA1* genes in maize, sorghum and setaria. We show that *RA1* expression marks boundary domains adjacent to meristems in sorghum and setaria inflorescences in concordance with *RA1* transcript accumulation in maize. We generated *RA1* transgenes from maize (*Zm*), sorghum (*Sb*) and setaria (*Sv*) loci and utilized the strong maize *ra1-R* mutant to investigate the impact of expressing *ZmRA1*, *SbRA1* and *SvRA1* transgenes on the regulation of branching in maize tassels and ears. Expression as a transgene of *ZmRA1* including flanking upstream and downstream sequences recovered normal inflorescence morphologies in *ra1-R* mutants. Interspecies expression of two transgene variants of the *SbRA1* locus, one modeled as the entire endogenous tandem duplication and the other as only the non-frameshifted downstream gene copy, yielded a range of *ra1-R* inflorescence architectures, showing partial recovery with or without novel branch patterns and fasciation. We found that interspecies expression of an *SvRA1* transgene, which lacks *cis*-promoter sequences conserved in maize, sorghum and other Andropogoneae species, either not at all or only partially recovered normal inflorescence forms in *ra1-R* mutants, whereas fusing the *SvRA1* coding region to the *ZmRA1* upstream region recovered normal inflorescence morphology in *ra1-R* mutants. Our functional tests of *RA1* sufficiency indicate that heterochronic modulation of meristem determinacy that results from *cis*-regulatory differences impacts ear and tassel morphology, and is a likely driver of inflorescence diversity throughout the grasses.

## RESULTS AND DISCUSSION

### Inflorescence architectures and *RA1* alleles in PACMAD and Panicoid grasses

Mature maize inflorescences are spatially and morphologically distinct and produce dimorphic, unisexual florets: a terminal tassel bearing staminate florets and a lateral ear with pistillate florets (**Fig. 1A, B**). Mutations in the *ZmRA1* gene, typified by the strong *ra1-R* allele (Vollbrecht et al., 2005), result in multiple orders of branching in the tassel and the ear (**Fig. 1C, D**) that resemble complexly branched inflorescences of other grasses, such as terminal panicles of sorghum (**Fig. 1E**) and setaria (**Fig. 1F**) which have unimorphic, bisexual florets. The conspicuous diversity of mature inflorescence morphologies in maize, sorghum and setaria, largely attributed to variation in degree of branching, manifests early in development (**Fig. S1**). Maize, sorghum and setaria belong to the subfamily Panicoideae, and within this large clade of grasses, maize and sorghum are members of tribe Andropogoneae, whereas setaria is a member of tribe Paniceae (ref. Kellogg, 2015). Maize and sorghum inflorescences produce a multitude of spikelets in pairs as is characteristic of related species in the Andropogoneae, whereas the setaria inflorescence is dense with single spikelets that each develop in close association with a bristle (Doust and Kellogg, 2002; Kellogg, 2015).

**Figure 1.**
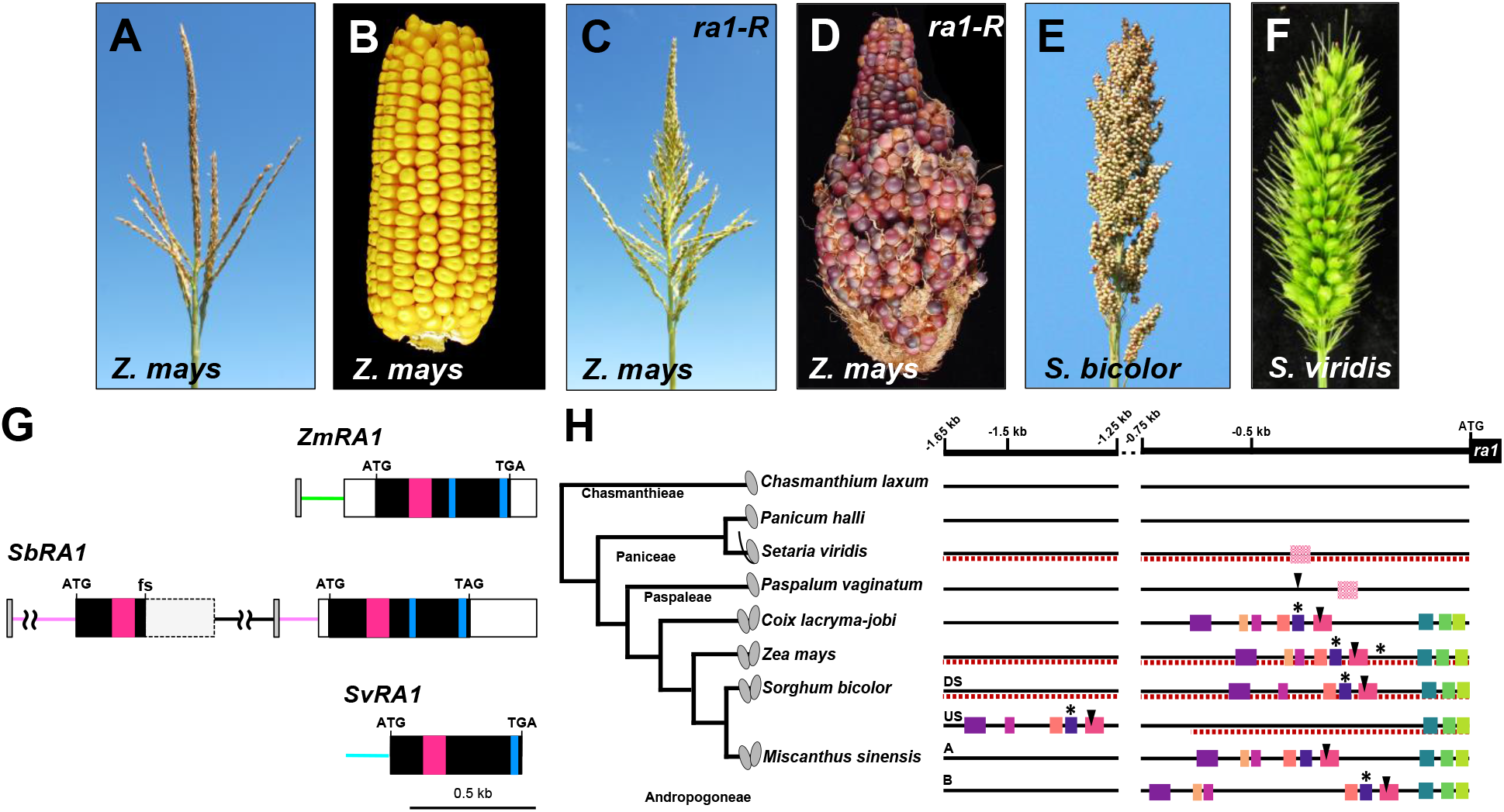
Architecture of maize, maize *ra1-R* mutant, sorghum and *S. viridis* inflorescences and genomic relationship of *RA1*. Normal inbred B73 maize tassel (A) and ear (B). Maize *ra1-R* mutant tassel (C) and ear (D). (E) *S. bicolor* panicle. (F) *S. viridis* panicle. (A-F) Inflorescences not to scale. (G) Annotated gene structure for *RA1* homologs. Tandem duplication of *SbRA1* locus is shown with indicated frameshift mutation (fs) in upstream copy of *SbRA1*. Predicted promoter regions are indicated by color lines. Gray box, conserved non-coding *cis* sequences (see 1H). Open box, UTR sequences. Magenta box, encoded C_2_H_2_ zinc finger domain. Blue box, encoded EAR motif. (H) Conserved non-coding *cis* sequences in the *RA1* promoters of Panicoid grasses. Among species in the tribe Andropogoneae, the promoter regions of *RA1* display different motifs conserved in sequence and arrangement (correspondingly colored boxes are conserved; Fig. S3C) compared to other tribes in the Panicoideae family. Upstream (US) and downstream (DS) tandem duplicate *SbRA1* copies and duplicate *MsRA1* copies A and B are indicated. Dashed lines underscore promoter regions incorporated into transgene cassettes. Some conserved sequences contained binding motifs for well-known transcriptional regulators, such as LEAFY and Clade A ARFs (Fig. S3B). Solid squares, *P*-values ≤ 1^-20^; cross-hatched squares, *P*-values ≤ 1^-05^; arrowhead – LEAFY-binding motifs; asterisks – Clade A ARF-binding motifs. Character state of spikelets (paired, single or with a bristle) is indicated on the phylogeny.

Comparative genomic data indicate the *RA1* locus is specific to the PACMAD clade, whose largest subfamilies are the Panicoideae and Chloridoideae, where the intronless structure and unique QGLGGH motif within the C_2_H_2_ zinc finger present in maize (Vollbrecht et al., 2005) appear conserved. For example, a syntenic copy of *RA1* is absent from the genomes of BOP clade members rice (*Oryza sativa*), *Brachypodium distachyon* and wheat (*Triticum aestivum*) (**Fig. S2A**) (Vollbrecht et al., 2005; Sigmon, 2010), but is present in the genome assemblies of Chloridoideae species teff (*Eragrostis tef*) and *Oropetium thomaeum* (Schnable, 2019), and of finger millet (*Eleusine coracana*). Within the Panicoideae *RA1* resides as a single copy gene in maize and setaria and as a single-locus tandem duplication in sorghum (*SbRA1^TAN^* comprised of *SbRA1* upstream [*SbRA1^US^*] and *SbRA1* downstream [*SbRA1^DS^*] copies); however, a frameshift mutation in *SbRA1^US^* introduces a stop codon after the C_2_H_2_ zinc finger domain, rendering it presumably nonfunctional (**Fig. 1G**) (Vollbrecht et al., 2005; Sigmon, 2010). Previously published RT-PCR and transcript profiling data indicate that *SbRA1^US^* is not expressed in inflorescences of sorghum BTx623, while *SbRA1^DS^* is (Vollbrecht et al., 2005; Wang et al., 2018; Leiboff and Hake, 2019). Broad sampling of diverse cultivated and wild sorghums found that, in all cultivated accessions, 1) *SbRA1^US^* contains the same frameshift and that the *SbRA1^DS^* open reading frame (ORF) encodes a predicted full length RA1 protein; 2) the *SbRA1* tandem duplication likely originated relatively recently with the *Sorghum* genus and may not be present in other grass species (Sigmon, 2010). Two *RA1* loci are present in miscanthus (**Fig. 1H**), but these are segmental duplicates in this paleotetraploid species (Sigmon, 2010; Mitros et al., 2020). The encoded *Sb*RA1^DS^ protein of cultivated sorghums, hereafter referred to as *Sb*RA1, is ∼69% identical to the *Zm*RA1 protein and ∼56% identical to the *Sv*RA1 protein. *Zm*RA1 and *Sv*RA1 proteins are ∼65% identical. *Zm*RA1, *Sb*RA1 and *Sv*RA1 proteins share a highly conserved C_2_H_2_ zinc-finger domain and a conserved C-terminal EAR motif (**Figs. 1G and S2B**). Biochemical experiments have demonstrated the C_2_H_2_ zinc-finger domain binds DNA (Dathan et al., 2002), and the EAR motif acts as a potent transcriptional repressor (Hiratsu et al., 2004; Tiwari et al., 2004). The motifs and their positioning are highly conserved between *Zm*RA1, *Sb*RA1 and *Sv*RA1 proteins. The C_2_H_2_ zinc-finger domain between *Zm*RA1 and *Sb*RA1 differs by one conservative amino acid variant (I67V, position relative to *Zm*RA1) that is identical (V) between *Sb*RA1 and *Sv*RA1. Relative to *Zm*RA1 and *Sb*RA1, the *Sv*RA1 zinc-finger domain differs at three positions, none of them among invariant core C_2_H_2_ residues (Vollbrecht et al., 2005). The C-terminal EAR motif is conserved between *Zm*RA1 and *Sb*RA1 and varies by one residue (Q169E) in *Sv*RA1. A second EAR motif adjacent to the C_2_H_2_ zinc-finger domain (Sigmon 2010, Gallavotti et al., 2010) is highly conserved between *Zm*RA1 and *Sb*RA1 but absent from *Sv*RA1 (**Figs. 1G and S2B**). Physical interaction between *Zm*RA1 and REL2 involves both EAR motifs (Gallavotti et al., 2010); however, functional sufficiency of the maize C-terminal EAR motif has not been demonstrated.

By mining two kilobases of the *RA1* promoter region from eight Panicoideae taxa across the Chasmanthieae, Paniceae, Paspaleae and Andropogoneae tribes, we identified several blocks of highly conserved, noncoding *cis* sequence restricted to the Andropogoneae, where spikelets are paired (**Figs. 1H and S3**). These conserved *cis* sequences located in the promoter region of *ZmRA1* and *SbRA1^DS^* (Sigmon, 2010), were absent from the ∼0.7 kb promoter region included in our *SbRA1^US^* transgene construct and were largely absent or not well conserved outside the Andropogoneae, including in *SvRA1* (**Figs. 1H and S3A, S3B**). Within the four Andropogoneae tribe taxa, where there are six promoter regions due to gene duplications, the conserved non-coding *cis* sequences harbored 48 putative transcription factor binding sites present among at least five of six sequences queried (**Supplemental dataset**). In maize, some of the conserved *cis* sequence overlaps with accessible chromatin profiled from developing ears but not accessible in leaves (**Fig. S3C**; Ricci et al., 2019). Indeed, coinciding with the region of accessible chromatin we found that DNA affinity purification (DAP) sequencing of maize AUXIN RESPONSE FACTOR (ARF) transcription factors identified binding peaks (**Fig. S3C**; Galli et al., 2018) centered on a putative ARF binding motif, providing a possible additional link between auxin signaling and response and branch development (Gallavotti et al., 2008; Eveland et al., 2014). Also within the region of accessible chromatin and within a conserved non-coding *cis* sequence we identified a putative LEAFY (LFY) transcription factor binding motif (Winter et al., 2011) in all six Andropogoneae sequences queried (**Figs. 1H, S3B, S3C and Supplemental dataset**). LFY is bifunctional as an activator and repressor in Arabidopsis (William et al., 2004; Winter et al., 2011). Within the Andropogoneae the protein-coding regions of the *LFY*-like genes are highly conserved suggesting purifying selection and constraint on amino acid sequence (Bomblies and Doebley, 2005). Interestingly, in maize, transcripts of the *LFY* homologs *Zea FLORICAULA/LEAFY1* (*ZFL1*) and *ZFL2* (Bomblies et al., 2003) accumulate in SPMs in a pattern that would likely border *ZmRA1* transcript accumulation (Vollbrecht et al., 2005). Tassel branch number is decreased in *zfl1; zfl2* double mutants, and positively correlates with *ZFL2* copy number (Bomblies et al., 2003; Bomblies and Doebley, 2006). These *ZFL* data are consistent with negative regulation of *ZmRA1* activity by *ZFL* gene activity, making it tempting to speculate that ZFL could repress *ZmRA1* where their expression domains abut in boundary cells at the margin of SPMs.

### *RA1* marks boundary domains adjacent to meristems in sorghum and setaria panicles

To determine the accumulation of *RA1* transcripts in sorghum and setaria inflorescences, we performed RNA *in situ* hybridization with an antisense probe for *ZmRA1*, along with the meristem marker gene *KNOTTED1* (*KN1*; Jackson et al., 1994). In sorghum, *RA1* transcripts accumulated in a boundary domain directly adjacent to the SPM, as marked by accumulation of *KN1* transcripts (**Figs. 2A, B** and **S4A, B**). *RA1* transcripts were not detected in early-staged setaria inflorescences initiating branch meristems, as shown by accumulation of *KN1* (**Fig. 2C, D**), consistent with transcriptomic profiling of setaria inflorescence development (Zhu et al., 2018). In later-staged setaria inflorescences marked by SMs and bristles, we detected *RA1* transcripts in accordance with transcriptomic data (Zhu et al., 2018), which showed boundary domain accumulation adjacent to the SM (**Figs. 2E, F** and **S4C-F**). We consistently did not detect *RA1* transcript accumulation in or adjacent to bristles, further distinguishing them from the spikelets they are paired with. In maize, *RA1* transcripts accumulate between recently-initiated SPMs and the inflorescence or branch axis (Vollbrecht et al., 2005). These results demonstrate 1) a conserved spatial pattern of *RA1* transcript accumulation that marks boundary domains adjacent to spikelet-associated short branch meristems in sorghum, setaria and maize inflorescences, whether SMs (setaria) or SPMs (maize and sorghum), 2) a conserved lack of expression associated with BMs and LBs and other branch types (i.e. the bristle in setaria) and 3) distinct temporal patterns consistent with discrete branching ontogenies.

**Figure 2.**
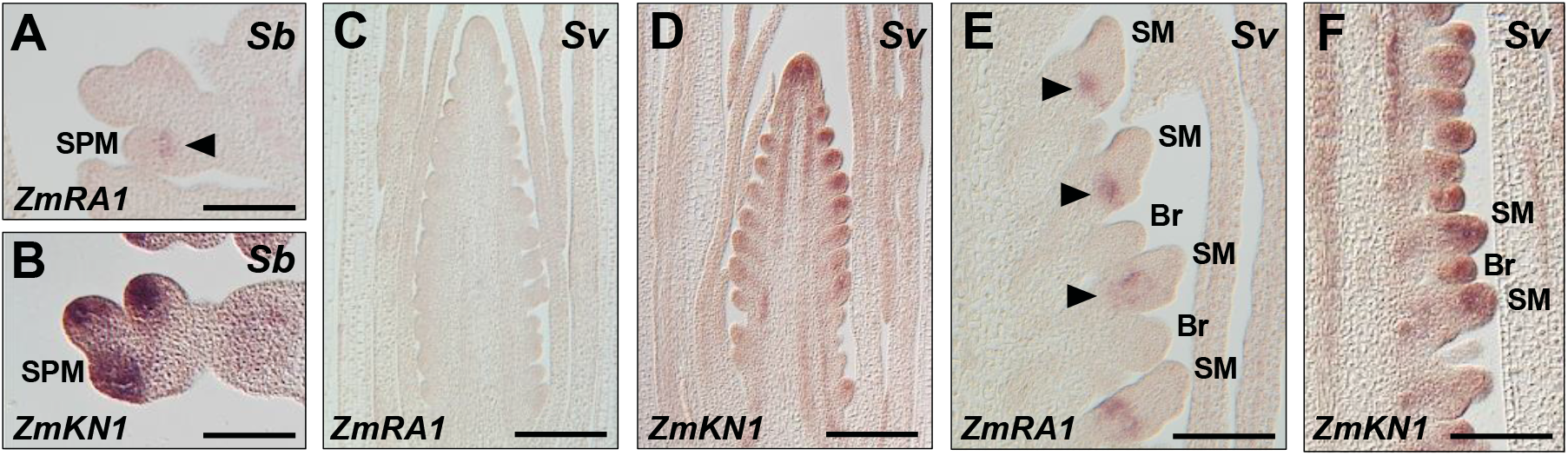
RNA *in situ* hybridization in sorghum and *S. viridis* inflorescences. Antisense RNA probes to *ZmRA1* (A, C, E) or *ZmKN1* (B, D, F) were hybridized to longitudinal sections of developing inflorescences from sorghum, *Sb* (A, B) or *S. viridis, Sv* (C-F). Arrowheads denote *RA1* transcript accumulation in boundary domains. Scale bars, 100 µm.

### Expression of a *ZmRA1* transgene largely recovers normal inflorescence architectures in *ra1-R* mutants

To study the function of promoter *cis* and coding sequence diversity of *RA1* loci in shaping the inflorescences of maize, sorghum and setaria, we generated a suite of transgenic experiments using interspecies gene transfer (Nikolov and Tsiantis, 2015). Maize, sorghum and setaria *RA1* genes and one chimeric maize-setaria *RA1* gene were introduced into maize and backcrossed into the B73 inbred genetic background containing the *ra1-R* mutant allele. During backcrosses the events were scored for evidence of a heritable, single-locus, herbicide resistance phenotype as an indicator of stable expression of the 35S::BAR component of the transgene cassette. In total 17 independent transgenic events satisfied these genetic segregation criteria (**Table S2**) and these were also scored qualitatively for their capacity to complement the *ra1-R* mutant phenotype; from among them, we selected nine events for detailed analysis (**Table S1 and Methods**).

To examine maize *RA1* gene function, we first asked if normal tassel and ear morphologies could be recovered in severe *ra1-R* mutants expressing a reintroduced *ZmRA1* genomic fragment containing 2.95 kb of the promoter region including the conserved *cis* sequences as well as 2.35 kb of sequence downstream of the CDS. We refer to this transgenic cassette as ‘*198*’ (**Fig. 3A; Table S1**). Five independent, stable, single-locus transgene events were generated for 198. Four of them showed similar effects on the *ra1-R* phenotype and minimal pleiotropy while the fifth was markedly pleiotropic (**Supplemental Table 1**), conferring a dwarfed plant stature and severely reduced tassels and ears. We studied the effects of *198* in a single, non-pleiotropic insertion event (**Table S2**). Gross tassel and ear morphology of *ra1-R* mutants expressing *198* appeared normal relative to non-transgenic *ra1-R* siblings (**cf. Figs. 3B-E to 1A**). Notably, *ra1-R* ears expressing *198* were fully unbranched, and kernels were in straight parallel rows along the ear axis; in contrast, kernel rowing was crooked in highly ramified *ra1-R* ears (**Fig. 3D, E**) (Vollbrecht et al., 2005).

**Figure 3.**
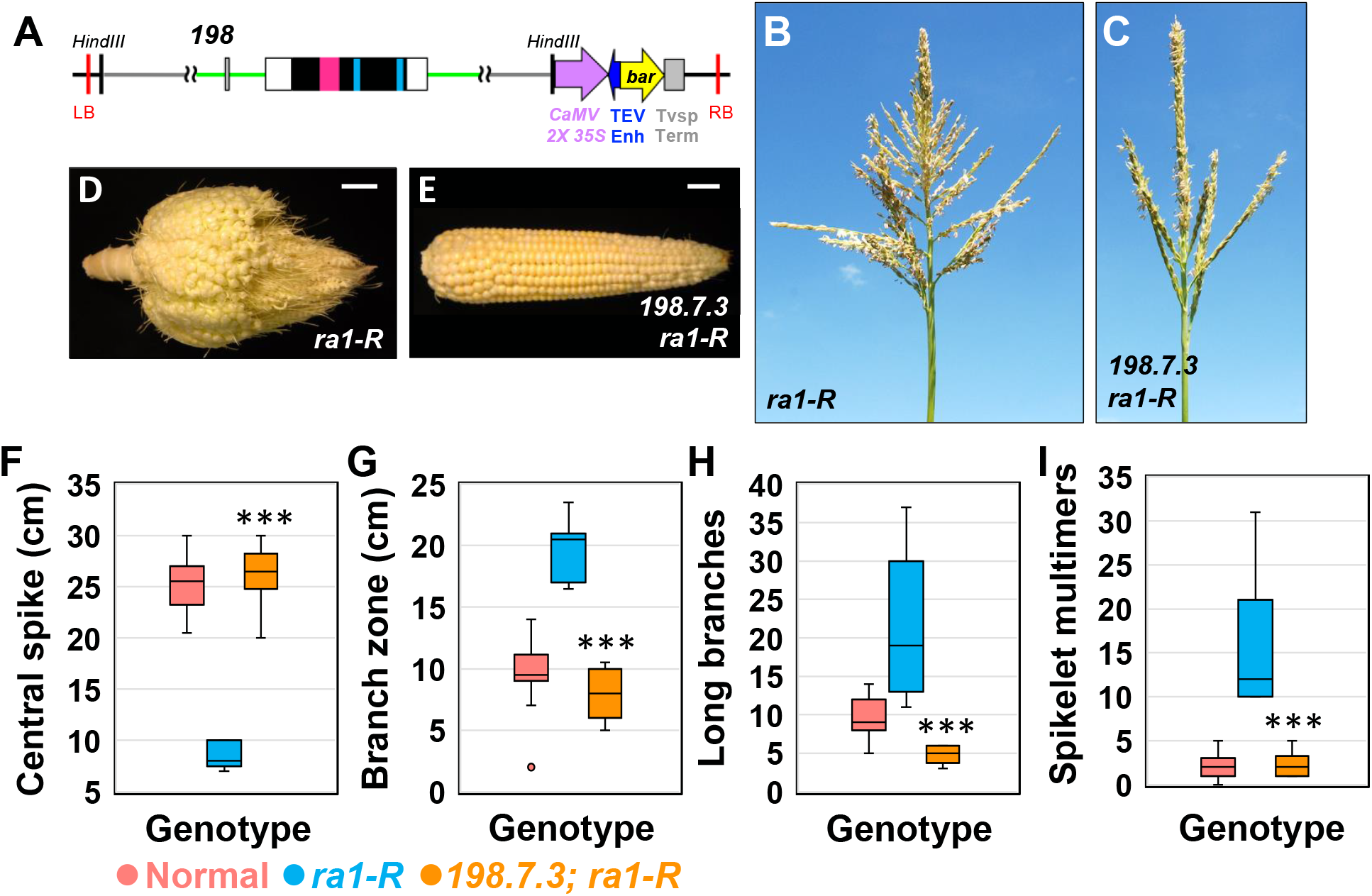
Expression of the *ZmRA1* locus as a transgene in *ra1-R* mutant background. (A) *198* cassette for expression of *ZmRA1* containing 2.9 kb of upstream sequence including conserved non-coding *cis* regions. (B) *ra1-R* tassel. (C) *ra1-R* tassel expressing 198.7.3. (D) *ra1-R* ear. (E) *ra1-R* ear expressing 198.7.3. Scale bars, 2 cm. (F) Central spike length. (G) Branch zone length. (H) Number of long branches. (I) Number of spikelet multimers. For all box and whisker plots, the bottom and top boxes represent the first and third quartile, respectively, the middle line is the median, and the whiskers represent the minimum and maximum values, outlier data points are displayed as individual dots. Two-tailed Student’s *t* test for transgene vs. *ra1-R* \*\*\**P*<0.001; normal, n = 20; *ra1-R*, n = 10; 198.7.3, n = 8.

We quantified degree of branching, including branch type, lengths and spikelet pair density (**Fig. S5**) among inflorescences of segregating normal, *ra1-R* mutants expressing *198* and non-transgenic *ra1-R* siblings to evaluate the degree of normal phenotype recovery. Along the primary axis of the tassel, normal maize produces LBs at the base with an immediate shift to short branches of SPs on the central spike (**Fig. 1A**). *ra1-R* mutants produce LBs at the tassel base, then a variable number of transformed, mixed-fate branches bearing both SPs and single spikelets, followed by transformed branches (“spikelet multimers”) with multiple, single spikelets and finally an abbreviated central spike predominantly of short branches of SPs (**Fig. 3B**) (Vollbrecht et al., 2005). The length of the central spike (CS) between normal and *ra1-R* expressing *198* were nearly equivalent (mean difference +0.95 cm); CS was significantly longer in *ra1-R* with the transgene compared to non-transgenic *ra1-R* siblings (mean difference -17.68 cm) (**Fig. 3F**). The length of the long branch zone (LBZ) was slightly shorter in *ra1-R* expressing *198* relative to normal (mean difference -1.68 cm), whereas LBZ was significantly shorter in transgene positive *ra1-R* compared to non-transgenic *ra1-R* siblings (mean difference -11.84 cm) (**Fig. 3G**). Normal tassels produced on average 4.9 more LBs compared with *ra1-R* tassels expressing *198*, whereas non-transgenic *ra1-R* siblings produced on average 17.4 more LBs than *ra1-R* expressing *198* (**Fig. 3H**). We observed a negligible difference in spikelet multimers (referred to as ‘multimers’ throughout) between normal and *ra1-R* transgene-expressing tassels, but non-transgenic *ra1-R* siblings produced on average 14 more multimers than *ra1-R* expressing *198* (**Fig. 3I**). Spikelet pair density (SPD) taken from a circumference of 1 cm at the CS midpoint was lower in *ra1-R* transgene positive plants compared with both normal and non-transgenic *ra1-R* siblings (-3.2 and -2.67 SPs, respectively) (**Fig. S6B**). The three most-basal tassel LBs were longer in *ra1-R* expressing *198* compared with both normal and non-transgenic *ra1-R* siblings (**Fig. S6C**). Collectively, these results indicate that the *ZmRA1* transgene is sufficient to recover normal inflorescence architectures in the *ra1-R* mutant background.

### Interspecies expression of a tandem duplicated *SbRA1* modeled transgene produces novel *ra1-R* inflorescence architectures

We next asked if interspecies expression of the canonical tandem duplicated *SbRA1* locus could recover normal tassel and ear morphologies in *ra1-R* mutants. We modeled the tandem duplicated *SbRA1* transgenic cassette *SbRA1^TAN^* as a 6 kb genomic DNA fragment that includes ∼0.7 kb promoter region of *SbRA1^US^*, the *SbRA1^US^* paralogous coding region followed by the contiguous 2.03 kb (including the conserved *cis* sequences) between the *SbRA1^US^* paralogous stop codon and the beginning of the *SbRA1^DS^* predicted ORF, the predicted ORF and 2.17 kb downstream of the *SbRA1^DS^* stop codon. We refer to this construct as ‘*195*’ (**Fig. 4A; Table S1**). Three independent, stable, single-locus transgene events were generated for *195* and backcrossed into the B73 background; we studied its effects on the *ra1-R* mutant in all three (**Table S2**).

**Figure 4.**
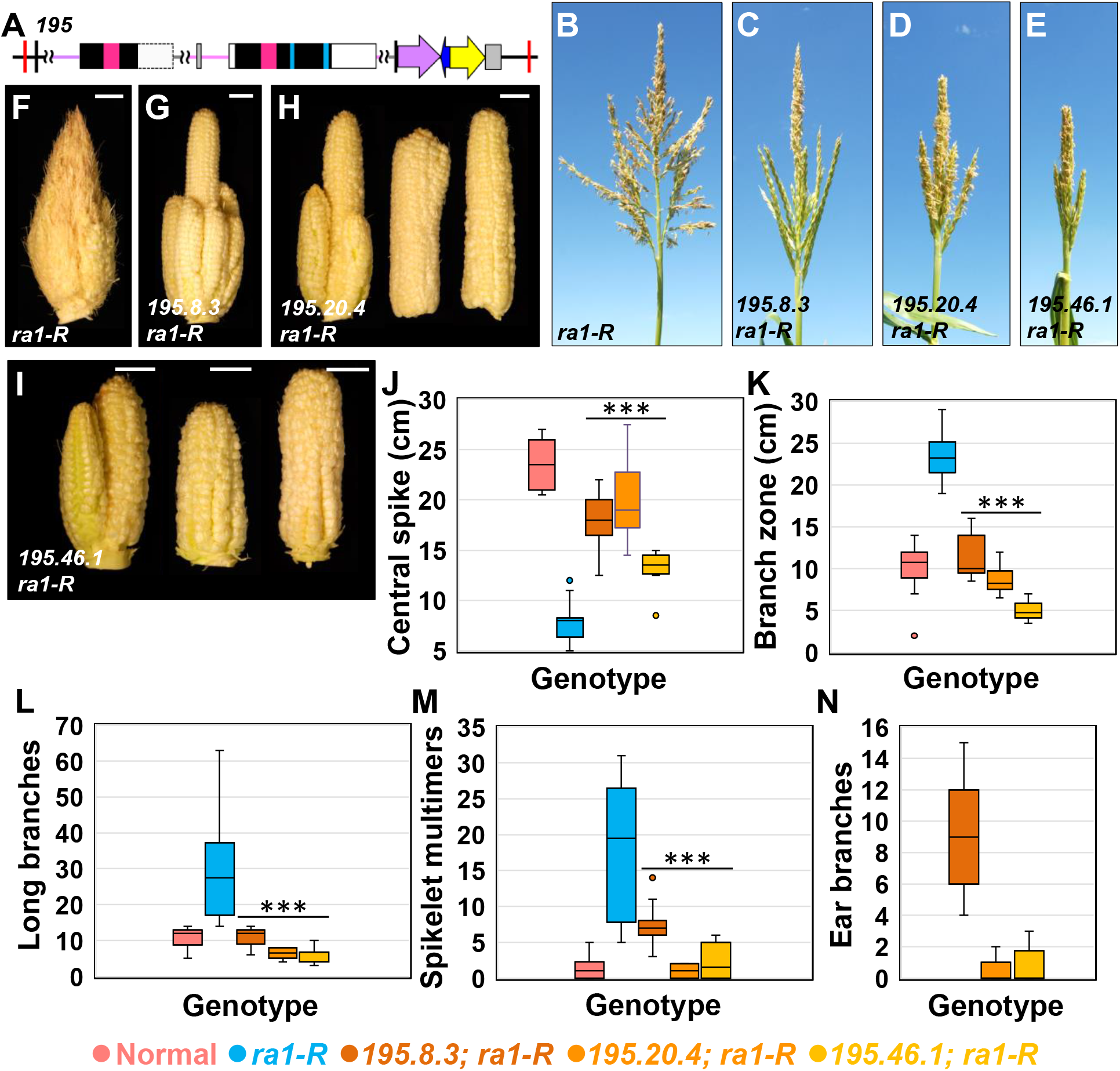
Interspecies expression of the tandem duplicated *SbRA1* modeled transgene in the *ra1-R* mutant background. (A) *195* cassette for interspecies expression of the tandem duplicated *SbRA1* locus. (B) *ra1-R* tassel. (C-E) *ra1-R* tassels expressing 195.8.3 (C), 195.20.4 and 195.46.1 (E) transgenes. (F) *ra1-R* ear. (G-I) *ra1-R* ears expressing 195.8.3 (G), 195.20.4 (H) and 195.46.1 (I) transgenes. Scale bars, 2 cm. (J) Central spike length. (K) Branch zone length. (L) Number of long branches. (M) Number of spikelet multimers. (N) Number of ear branches. For all box and whisker plots, the bottom and top boxes represent the first and third quartile, respectively, the middle line is the median, and the whiskers represent the minimum and maximum values, outlier data points are displayed as individual dots. Two-tailed Student’s *t* test for transgene vs. *ra1-R* \*\*\**P*<0.001; *ra1-R*, n = 18; 195.8.3, n = 11; 195.20.4, n = 12; 195.46.1, n = 11.

Overall, tassels of *ra1-R* mutants that expressed *195* were much less branched and ranged from normal (events 195.8.3 and 195.20.4) to compact (event 195.46.1) relative to highly branched non-transgenic *ra1-R* siblings (**Fig. 4B-E**). Similarly, *195*-expressing *ra1-R* ears displayed a range in gross phenotype (**Fig. 4F-I, N)**, but were overall much less branched than *ra1-R* sibling ears. For event 195.8.3, ear branching was reminiscent of weak *ra1* mutant alleles (**Fig. 4G**) (Vollbrecht et al., 2005; Gallavotti et al., 2010). Ears from event 195.20.4 and 195.46.1 were occasionally fasciated and branched, and frequently had crooked kernel rows (**Fig. 4H, I, N**). Ears from event 195.46.1 were consistently short and compact (**Fig. 4I**).

To understand the impact of *195* on *ra1-R* inflorescences, we quantified branch phenotypes for the three events. When compared to non-transgenic *ra1-R* siblings, mean CS lengths were significantly longer (range of differences from +5.61 to +12.15 cm) and mean LBZ lengths were significantly shorter in *ra1-R* carrying the *195* transgene (range of differences from -12.13 to -18.36 cm) (**Fig. 4J, K**). Non-transgenic *ra1-R* siblings produced on average 29.11 LBs and 18 multimers, which was significantly more compared to the mean range of 5.17 to 10.73 LBs and 2.33 to 7.09 multimers in *195* expressing *ra1-R* siblings (**Fig. 4L, M**). SPD had a mean range of differences from -0.25 to +5.5 SPs between *ra1-R* expressing the *195* transgene and non-transgenic *ra1-R* siblings (**Fig. S7B**). The three most basal LBs were significantly shorter in *ra1-R* tassels that expressed the *195* transgene compared with non-transgenic *ra1-R* siblings (**Fig. S7C**).

Long branches are completely suppressed in normal ears (**Fig. 1B**); LBs are de-repressed by mutations in *ZmRA1* (**Fig. 1D**) (Vollbrecht et al., 2005). Ears of strong *ra1* mutant alleles, such as *ra1-RSd*, produce over 200 branches (Weeks, 2013). *ra1-R* ears expressing the *195* transgene were significantly less branched compared to highly branched ears of non-transgenic *ra1-R* siblings (**Fig. 4N**). Event 195.8.3 had a mean ear branch number of 9.3, similar to previously reported mean ear branch totals for weak alleles, *ra1-63.3359* (11.2 branches) or *ra1-RS* (12.1 branches) (Weeks, 2013). Events 195.20.4 and 195.46.1 had a mean of <1 branch (**Fig. 4N**). Transcripts of the *195* transgene accumulated in developing tassels beyond the stages when the endogenous *ZmRA1* transcript accumulation are highest (**Fig. S8A**), supporting heterochronic expression of the transgene in the tassel.

Taken together, expression of the *195* transgene reduced the order of branching in *ra1-R* mutant inflorescences, but curiously also produced novel *ra1-R* phenotypes that included compact tassels and ears, and ear fasciation (**Fig. 4D, E, H, I**). Pleiotropic fasciation and stubbiness in the main axis suggest effects on the main inflorescence meristem, where *ra1* expression was not detected in normal maize or sorghum. Strong, likely null, maize *ra1* alleles have genetic lesions in the C_2_H_2_ zinc finger domain (Vollbrecht et al., 2005), a putative DNA binding domain (Dathan et al., 2002). Indeed, *Zm*RA1 is suggested to bind and modulate the expression of hundreds of genes during tassel and ear development, which includes the putative direct targeting and repression of *COMPACT PLANT2* (*CT2*; Bommert et al., 2013; Eveland et al., 2014). Loss-of-function *ct2* mutants have compact inflorescences and fasciated ears (Bommert et al., 2013), similar to what was observed to be conditioned by the *195* transgene (**Fig. 4B-I**). To explain the novel *ra1-R* phenotypes, we hypothesize that the *195* transgene may function ectopically and affect expression of target genes like *CT2* outside of the spatio-temporally normal expression domain for *RA1*. Misregulation of *RA1* could occur if the upstream copy competes with the downstream copy for binding of regulatory factors, or if the gene duplication itself alters regulation, for example, by changing the distance between *cis*-regulatory elements or by creating novel ones. Another potential mechanism for the novel phenotypes could be at the level of the gene product. For example, given that the truncated upstream *RA1* copy encodes a C_2_H_2_ zinc finger domain (**Figs. 1G** and **4A**), expression from both copies could lead to binding interference between *Sb*RA^US^ (truncated) and *Sb*RA^DS^ (complete) proteins, where *Sb*RA^DS^ is required at sufficient levels to impose meristem determinacy. Similar interference mechanisms for dominant negative alleles have been reported to influence flowering in Arabidopsis (Ahn et al., 2006) and sunflower (Blackman et al., 2010). Although *SbRA^US^* expression is barely detectable in sorghum inflorescences (Vollbrecht et al., 2005), we did not assay its expression in the transgenic lines.

### Interspecies expression of the downstream *SbRA1* modeled transgene partially recovers normal inflorescence architectures in *ra1-R* mutants

Because *195* conditioned novel phenotypic changes in addition to complementation, we asked if normal tassel and ear morphologies in *ra1-R* mutants could be recovered by interspecies expression of only the downstream *SbRA1* locus, which does not contain frameshifts or apparent deleterious mutations. The downstream *SbRA1* transgenic cassette *SbRA1^DS^* was modeled to include its predicted ORF and 1.68 kb upstream including the conserved *cis* sequences plus 2.17 kb downstream of the stop codon, and we refer to this construct as ‘*196*’ (**Fig. 5A; Table S1**). Three independent, stable, single-locus transgene events were generated for *196* and backcrossed to the *ra1-R* mutant in B73, and we studied its effects in all three (**Table S2**).

**Figure 5.**
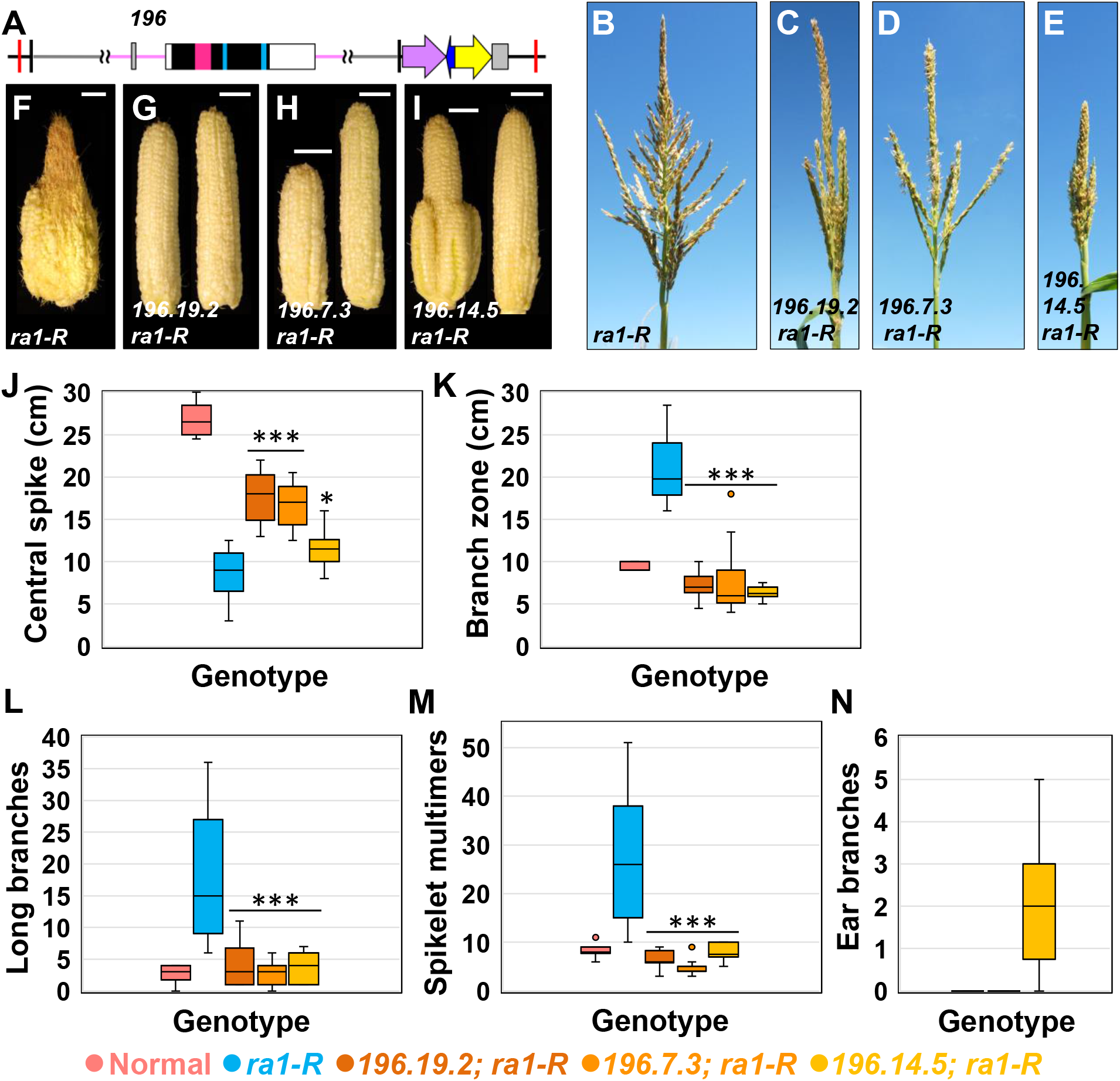
Interspecies expression of the downstream *SbRA1* modeled transgene in the *ra1-R* mutant background. (A) *196* cassette for interspecies expression of the downstream *SbRA1* locus. (B) *ra1-R* tassel. (C-E) *ra1-R* tassels expressing 196.19.2 (C), 196.7.3 (D) and 196.14.5 (E) transgenes. (F) *ra1-R* ear. (G-I) *ra1-R* ears expressing 196.19.2 (G), 196.7.3 (H) and 196.14.5 (I) transgenes. Scale bars, 2 cm. (J) Central spike length. (K) Branch zone length. (L) Number of long branches. (M) Number of spikelet multimers. (N) Number of ear branches. For all box and whisker plots, the bottom and top boxes represent the first and third quartile, respectively, the middle line is the median, and the whiskers represent the minimum and maximum values, outlier data points are displayed as individual dots. Two-tailed Student’s *t* test for transgene vs. *ra1-R* \*\*\**P*<0.001, \**P*<0.05; *ra1-R*, n = 15; 196.19.2, n = 10; 196.7.3, n = 12; 196.14.5, n = 10.

Overall, tassels from *ra1-R* mutants that expressed *196* were less branched and ranged from normal (events 196.19.2 and 196.7.3) to moderately compact (event 196.14.5) architectures relative to highly ramified architecture of non-transgenic *ra1-R* siblings (**Fig. 5B-E**). Similarly, *ra1-R* ears expressing the *196* transgene displayed a range in gross phenotype (**Fig. 5F-I, N)**. Events 196.19.2 and 196.7.3 produced unbranched ears with straight rows of kernels along the ear axis (**Fig. 5G, H**), whereas event 196.14.5 showed ear branching reminiscent of weak *ra1* mutant alleles (**Fig. 5I**) (Vollbrecht et al., 2005; Gallavotti et al., 2010).

To characterize the impact of *196* on *ra1-R* inflorescences in detail, we quantified tassel branch phenotypes for the three events. When compared to non-transgenic *ra1-R* siblings, mean CS lengths were significantly longer (range of differences from +2.75 to +8.95 cm) and mean LBZ lengths were significantly shorter in *ra1-R* that carried the *196* transgene (range of differences from -12.71 to -14.15 cm) (**Fig. 5J, K**). Non-transgenic *ra1-R* siblings produced on average 26.8 LBs and 17 multimers, which was significantly more compared to the mean range of 4.58 to 8 LBs and 2.83 to 4.2 multimers in *ra1-R* expressing the *196* transgene (**Fig. 5L, M**). SPD had a mean range of differences from +1.53 to +7.33 SPs between *ra1-R* with the *196* transgene and transgene-free *ra1-R* siblings (**Fig. S9B**). The three most basal LBs were significantly shorter in *ra1-R* tassels with the *196* transgene compared with non-transgenic *ra1-R* siblings (**Fig. S9C**). Interspecies expression of *196* was sufficient to impose SPM determinacy in *ra1-R* ears for events 196.19.2 and 196.7.3, where branch suppression was fully penetrant. Event 196.14.5 had on average 2 branches (**Fig. 5N**), which was significantly less than average ear branch number for weak *ra1* alleles (Weeks, 2013). Transcripts of the *196* transgene accumulated in developing tassels beyond the stages when the endogenous *ZmRA1* transcript accumulation are highest (**Fig. S8B**), supporting heterochronic expression of the transgene in the tassel.

Collectively, interspecies expression of *196* restored more normal ear inflorescences with less branching and straighter rows, and less pleiotropy with respect to fasciation and shortened axes, relative to the *195* cassette. Furthermore, both the *195* and *196* constructs substantially remediated *ra1-R* tassel branching. Given that the *196* transgene eliminates the *SbRA1^US^* locus present in the *195* construct, these results suggest functional *cis*-regulatory element(s) that reside in the 1.68 kb sequence promoter region of the *SbRA1^DS^* locus are affected by their proximity to *SbRA1^US^* in the tandem duplication, especially in the maize ear. Our data on *ra1-R* mutants expressing either *195* or *196* cassettes are consistent with a hypothesis raised previously (Vollbrecht et al., 2005): variation in inflorescence architecture, and thus degrees of determinacy, is attributed to the developmental timing of *RA1* expression and its activity, as reflected in the range of branch types observed among maize mutant alleles, transgene versions, or genetic diversity of *RA1* in maize and other grasses. Furthermore, these results suggest that developmental context of *RA1* activity in the tassel and ear is crucial in regulating determinacy (cf., ear and tassel phenotypes in **Figs. 4 and 5**). Indeed, quantification of ear and tassel branch number in the F_1_ hybrid generation of B73 x Mo17 introgressions homozygous for the weak allele *ra1-63.3359* showed additive effects on ear branching and over-dominance effects on tassel branching (Weeks, 2013).

### Interspecies expression of *SvRA1* only recovers near normal inflorescence branching in *ra1-R* mutants when chimeric with the *ZmRA1* promoter region

Given the complex genetic nature of the *SbRA1* locus, we sought to explore the impact of the single copy *SvRA1* on inflorescence morphology. We were also interested in testing the impact of the *cis* sequences found in promoter regions of *ZmRA1* and *SbRA1^DS^* and conserved among Andropogoneae grasses, as well as sufficiency of the single EAR motif in *Sv*RA1. We therefore compared and contrasted interspecies expression of the *SvRA1* coding region with its endogenous promoter region that largely lacks the conserved *cis* sequences with expression of the *SvRA1* gene body in *cis* with the maize promoter region (*pZmRA1*). We modeled the *SvRA1* transgene cassette to include 1.53 kb of the predicted *SvRA1* promoter region, the coding region and 1.97 kb downstream of the stop codon and we refer to it as ‘*162*’ hereafter (**Fig. 6A; Table S1**).

**Figure 6.**
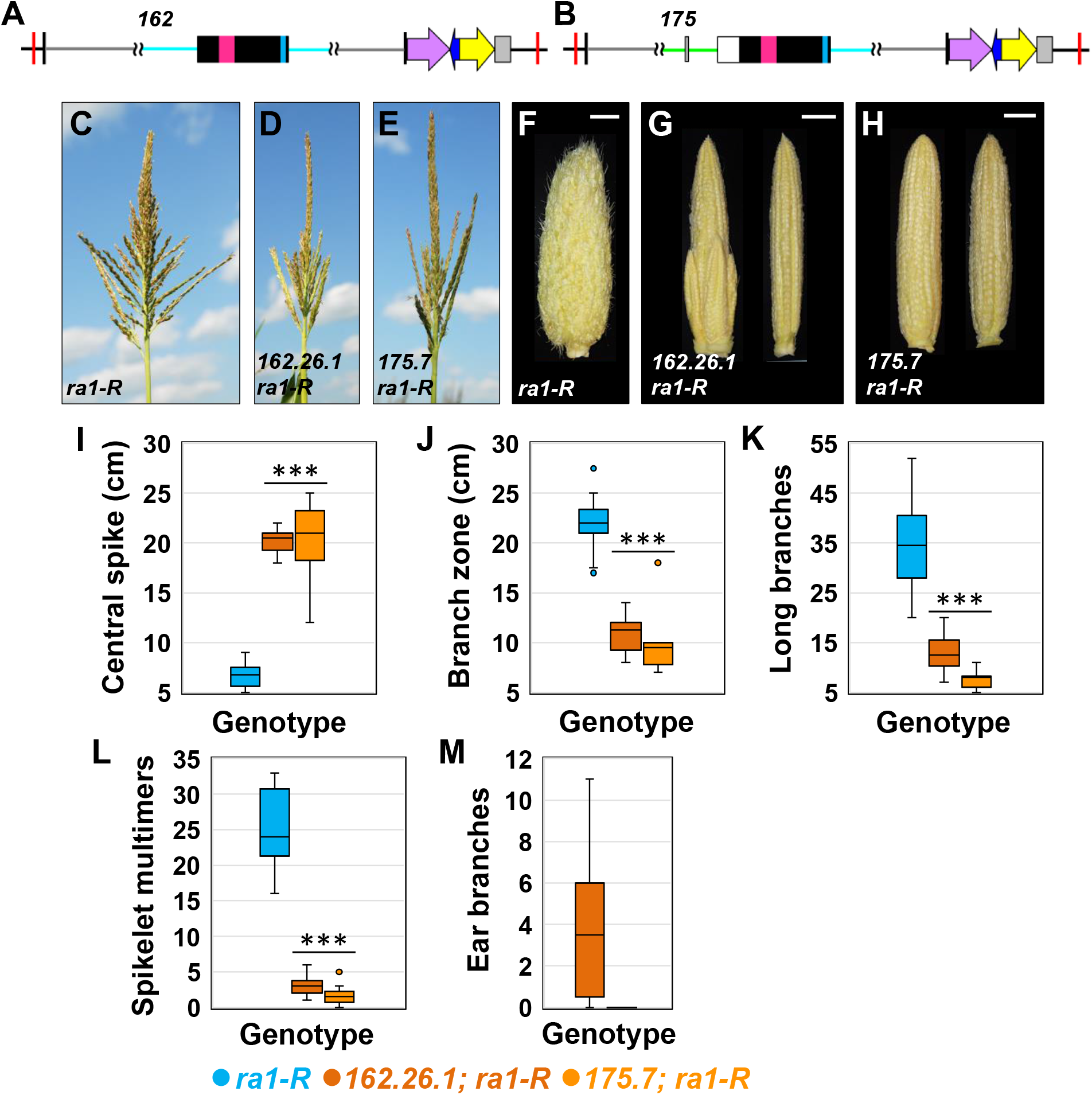
Interspecies expression of *SvRA1* or chimeric *SvRA1* as a transgene in the *ra1-R* mutant background. (A) *162* cassette for interspecies expression of the *SvRA1* locus. (B) *175* cassette for expression of the *SvRA1* coding region fused to the 2.9 kb *Zm* upstream region including conserved non-coding *cis* sequences. (C) *ra1-R* tassel. (D, E) *ra1-R* tassels expressing 162.26.1 (D) and 175.7 (E) transgenes. (F) *ra1-R* ear. (G, H) *ra1-R* ears expressing 162.26.1 (G) and 175.7 (H) transgenes. Scale bars, 2 cm. (I) Central spike length. (J) Branch zone length. (K) Number of long branches. (L) Number of spikelet multimers. (M) Number of ear branches. For all box and whisker plots, the bottom and top boxes represent the first and third quartile, respectively, the middle line is the median, and the whiskers represent the minimum and maximum values, outlier data points are displayed as individual dots. Two-tailed Student’s *t* test for transgene vs. *ra1-R* \*\*\**P*<0.001; *ra1-R*, n = 15; 162.26.1, n = 12; 175.7, n = 9.

Additionally, we generated a chimeric gene cassette termed *pZmRA1::SvRA1* where 2.95 kb of *ZmRA1* promoter region and five-prime untranslated region was fused upstream of the *SvRA1* coding sequence and 1.97 kb of downstream *SvRA1* sequence and we refer to the construct as ‘*175*’ hereafter (**Fig. 6B; Table S1**). Four independent, stably herbicide-resistant and single-locus transgene events were identified for *162* during backcrossing to the B73 tester line (**Table S2**). Of those events, three were unique among all stable, herbicide-resistant transgenics we propagated in this study, across all five constructs, in that they showed no notable effect on the strong *ra1-R* mutant phenotype or any other plant phenotypes examined. Thus, quantitative phenotyping was not performed for these three events, which strongly suggests the *SvRA1* transgene has little or no functional activity in maize. The fourth event for *162* showed some reduction of vegetative shoot stature and effects on inflorescence branching and was therefore examined for ear and tassel phenotype, although we consider it an outlier or novel event among the four *162* transgenic lines. One stable, single-locus transgene event was generated for *175* and backcrossed to B73 and it affected inflorescences but was non-pleiotropic for vegetative plant characteristics. Thus, we studied the effects of *162* and *175* in single-locus events backcrossed in the *ra1-R* mutant background (**Table S2**).

Overall, tassels from *ra1-R* mutants that expressed the novel *162* event or expressed the *175* transgene were less branched and had normal architectures relative to the highly branched architecture of non-transgenic *ra1-R* siblings (**Fig. 6C-E**). Similarly, *ra1-R* ears expressing *162* displayed a range in gross phenotype from unbranched ears with straight rows of kernels along the ear axis and no branches to those with crooked rows and a low degree of branching (**Fig. 6F, G, M**). In contrast, *ra1-R* ears expressing *175* were fully unbranched with kernels in straight parallel rows along the ear axis (**Fig. 6H, M**).

To understand the impact of the novel *162* event or of *175* on *ra1-R* tassels and ears, we quantified branch phenotypes. When compared to non-transgenic *ra1-R* siblings, mean CS lengths were significantly longer (difference +13.67 cm for both *162* and *175*) and mean LBZ lengths were significantly shorter in *ra1-R* tassels expressing either *162* or *175* transgenes (difference -11 cm for *162* and -12.1 cm for *175*) (**Fig. 6I, J**). Relative to normal tassels, mean CS lengths were shorter (difference -5.0 cm for both *162* and *175*) and mean LBZ lengths were marginally longer in *ra1-R* tassels with either *162* or *175* (difference +1.27 cm for *162* and +0.17 cm for *175*) (**cf. Figs. 3F, G to 6I, J**). Non-transgenic *ra1-R* sibling tassels produced on average 35.1 LBs and 25.1 multimers, which were significantly more compared to averages of 12.7 LBs and 3.1 multimers for *162*, and 7.5 LBs and 1.7 multimers for *175* expressing *ra1-R* siblings (**Fig. 6K, L**). Compared to a mean of 9.6 LBs and 2 multimers for normal tassels, *162* expressing *ra1-R* tassels produced on average 3.1 more LBs and 0.9 more multimers, whereas *175* expressing *ra1-R* tassels had 2.1 fewer LBs and 0.3 fewer multimers (**cf. Figs. 3H, I to 6K, L**). For SPD, *ra1-R* tassels with *162* had on average 3.2 more SPs along the CS compared to non-transgenic *ra1-R* siblings, and similarly, *ra1-R* tassels with *175* had 2.2 more SPs (**Fig. S10B**). Relative to SPD for normal tassels, *ra1-R* expressing *162* had on average 0.7 fewer SPs and *ra1-R* expressing *175* had 1.7 fewer SPs along the CS (**cf. Figs. S6B to S10B**). The three most basal LBs were consistently shorter in *ra1-R* tassels that carried the *162* transgene compared with non-transgenic *ra1-R* siblings; LBs were of similar length between *ra1-R* expressing the *175* transgene and non-transgenic *ra1-R* siblings (**Fig. S10C**). Compared to normal tassels, the three most basal LBs of *ra1-R* tassels expressing either *162* or *175* were shorter (**cf. Figs. S6C to S10C**).

Establishment of SPM determinacy during ear development differed conspicuously between *ra1-R* expressing the *175* transgene and expressing the novel *162* event. *ra1-R* with the *162* transgene produced an average of 4 branches, whereas *ra1-R* ears carrying the *175* transgene were unbranched (**Fig. 6M**). Overall, the *175* transgene behaved most similarly to the *198* endogenous maize construct.

Collectively, the transgene constructs containing *SvRA1* conferred degrees of complementation from non-to partial to nearly complete, all without inducing the novel inflorescence phenotypes of sorghum transgenes. Whereas in most *SvRA1* (*162*) lines the intact *SvRA1* gene did not complement the *ra1-R* mutant phenotype, we saw some effects in one line. Similarly, the *ZmRa1* and *SbRa1* events were not all identical in their phenotypic effects, as is not unusual among transgene events integrated into different chromosomal regions. We speculate that the novel *SvRA1* (*162*) event may be integrated in a genomic context that results in effectively ectopic expression, and therefore suggesting a lack of appropriate *cis* regulatory components in the *SvRA1* promoter region while revealing some functional potential of the SvRA1 gene product.

In the encoded polypeptides, *Zm*RA1 and *Sv*RA1 C_2_H_2_ zinc-finger domains vary by three amino acid residues, and the C-terminal EAR motif in *Sv*RA1 differs by one residue. However, a conserved EAR motif adjacent to the C_2_H_2_ zinc-finger domain in *Zm*RA1 is absent in *Sv*RA1 (**Figs. 1G and S2B**). In maize, RA1 physically interacts with REL2 via EAR motifs in a large transcriptional repressor complex to impose SPM determinacy (Gallavotti et al., 2010; Liu et al., 2019). Functional importance of the EAR motif adjacent to the C_2_H_2_ zinc-finger domain has not been tested genetically. Our data from the *175* chimeric gene cassette suggest the C_2_H_2_ -proximal EAR motif, which is by definition dispensable for RA1 function in setaria, is likewise to a significant degree nonessential in maize. Whereas complementation was only partial for novel *162* event, it was more complete for the chimeric *175* construct. The promoter region swap data clearly indicate that *cis*-encoded regulation of *RA1* expression is a key functional component in promoting SPM determinacy, especially during ear development. In an evolutionary context it is interesting to note that while spikelets are normally unpaired in setaria and the *SvRA1* gene is insufficient to complement the maize *ra1-R* mutant with its many unpaired spikelets, under the proper expression conditions the SvRA1 gene product does confer sufficient determinacy activity to restore SPs to *ra1-R* maize. These results suggest that within the Panicoideae subfamily of the PACMAD grasses *RA1* has an evolutionarily conserved determinacy function that contributes to specifying short branch meristems: SMs in setaria and SPMs in maize and sorghum. Our data are all consistent with a hypothesis wherein within the paired-spikelet Andropogoneae tribe, *RA1* has been adopted a key role in producing the SP by imposing determinacy in the proper developmental context rather than by specifying any strict SPM identity. It would be interesting to test whether the *RA1* genes from other Panicoid species as well as from Chloridoid subfamily and/or other PACMAD grasses show similar functions.

The developmental context in which genes and networks operate within meristems and flanking organ boundary domains is critical in determining inflorescence form. Elegant genetic studies on the spatiotemporal regulation and function of transcription factors have shed important light on the mechanisms governing inflorescence branching patterns. Genetic variation in distal regulatory elements (Clark et al., 2006; Studer et al., 2011), proximal or intronic *cis* regulatory elements (Arnaud et al., 2011; Wills et al., 2013; Kusters et al., 2015), coding sequences that alter protein function (Wang et al., 2005; Whipple et al., 2010), protein-protein interactions (Bartlett et al., 2016; Abraham-Juarez et al., 2020) or protein-DNA interactions (Maizel et al., 2005; Sayou et al., 2014) are critical drivers of inflorescence branching. Our data leveraging interspecies gene transfer and chimeric transgene expression suggest that *cis*-encoded regulation of *RA1* expression is a key factor in modulating meristem determinacy that ultimately impacts grass inflorescence architecture. With the ability to map hundreds of regulatory regions and transcription factor binding sites across diverse plant genomes (Lu et al., 2019; Galli et al., 2020), it will be important to understand the regulatory context of the conserved *cis* sequences that reside in *RA1* promoters.

Branch determinacy in the grasses is controlled by gene networks that function in boundary domains adjacent to the meristem they positionally regulate. Since their discovery, such ‘signaling centers’ have emerged as a major theme in regulating meristem determinacy, not meristem identity, and are key drivers of complex branching patterns seen in grass inflorescences (Whipple, 2017). Maize *RAMOSA* genes—*RA1* (Vollbrecht et al., 2005), *RA2* that encodes a LATERAL ORGAN BOUNDARY domain transcription factor (Bortiri et al., 2006) and the TREHALOSE PHOSPHATE PHOSPHATASE-encoding *RA3* (Satoh-Nagasawa et al., 2006)— constitute a ‘signaling center’ as these genes are co-expressed in overlapping boundary domains (Vollbrecht and Schmidt, 2009) and likely regulate a mobile signal that promotes determinacy of adjacent BMs. Similarly, BM determinacy is controlled by the GATA domain zinc-finger and SQUAMOSA PROMOTER BINDING PROTEIN transcription factors encoded by *TASSEL SHEATH1* (*TSH1*) and *TSH4* (Whipple et al., 2010; Chuck et al., 2010), and SM identity and determinacy are regulated by boundary expression of *BRANCHED SILKLESS1* and *INDETERMINANT SPIKELET1* that encode APETALA2 domain transcription factors (Chuck et al., 1998; 2007). BMs, SPMs and SMs are not meristem types found in eudicot inflorescences, where variation and complexity are largely governed by shifts in meristem identity (Prusinkiewicz et al., 2007; Lemmon et al., 2016). Given that *RA1* transcripts accumulate in meristem boundary regions during development of sorghum and setaria inflorescences, it will be interesting to test the functional consequences of mutating *RA1* in these grasses. Meristem identity genes in eudicots are expressed in meristems; genes that regulate inflorescence variation and complexity in the grasses are expressed in adjacent boundary domains to regulate meristem determinacy. Our work on the expression and functional conservation of syntenic *RA1* orthologs provides comparative insight into the genetic basis of grass inflorescence diversity, and opens the door for future reverse engineering of grass inflorescence evolution for crop improvement.

## MATERIALS AND METHODS

### Genetic Stocks

This study utilized the *ra1-R* allele (Vollbrecht et al., 2005) backcrossed seven generations to the B73 background to generate the “recurrent B73 parent;” either *ra1-R* homozygotes or *ra1-R/ra1-B73* heterozygotes were used in crossing schemes.

### Generation of *RA1* transgenes

The 35SBAR fragment from pTF101.1 was modified by PCR to introduce a *Hin*dIII site at the 3’ end of the terminator. This allowed a 2.0 kb *Hin*dIII restriction fragment containing 35SBAR-terminator to be isolated, treated with DNA polymerase I (Klenow) and dNTPs to generate blunt ends, and ligated into the *Sma*I site of pSB11 (Komari et al., 2006), creating a vector called pSB11_BAR. This vector, which contains the 35SBAR gene adjacent to and transcribed towards the T-DNA left border, was the precursor to all of the complementation vectors containing the genomic regions described below.

For construct *198*, *ZmRA1* and flanking regulatory regions were PCR amplified from *Z. mays* B73 genomic DNA and ligated with pSB11_BAR at *Hin*dIII. To distinguish the *198_RA1* allele from endogenous allele in subsequent generations after plant transformation, we introduced an *Acc*I restriction site in the *RA1* coding DNA sequence. This synonymous SNP (B73_v5 7: 114959005 C>T) is a natural, low frequency variant found in the maize inbred P39 haplotype (Vollbrecht et al., 2005). For the Sorghum construct *196*, a 6.0 kb *Xba*I fragment obtained by screening a BTx623-derived BAC library with a *ZmRA1* probe was cloned into pBluescript II KS (Agilent) and the *Hin*dIII site in the polylinker was used for ligation into pSB11_BAR. Sorghum construct *195* was generated from *196* following introduction of a *Hin*dIII site at the 3’ end of the upstream *SbRA1* frameshift copy (2: 58699332; **Table S1**) thereby removing a 1.6 kb fragment containing the upstream copy. For the *Setaria viridis* construct *162*, in-fusion cloning methods (Clontech/Takara) were employed to PCR-amplify and clone from *S.viridis* A10 genomic DNA a 4 kb fragment containing the *SvRA1* transcribed region and regulatory sequences into pSB11-BAR as a *Hin*dIII-*Bam*HI insertion. The maize/setaria chimeric construct *175* was generated as a translational fusion at the start codon by replacing the *Setaria* promoter-containing fragment in *162* with the 2.9 kb maize fragment. The reference genome coordinates of the *RA1* genes and regulatory regions are listed in **Table S1**, and all primers used for vector construction are listed in **Table S3**.

Constructs except for 175 were recombined into the pSB1 superbinary vector in *Agrobacterium tumefaciens* LBA4404 via triparental mating (Komari et al., 2006). These strains were used for Agrobacterium-mediated maize transformation of Hi-II embryos by the Iowa State University Plant Transformation Facility. Transgenic maize plants containing the *175* cassette were generated in Erik Vollbrecht’s lab at Iowa State University using particle bombardment of immature Hi-II embryos with the SB11_BAR-derived vector directly (Frame et al., 2000).

### Tests for recovery in *ra1-R*

T0 transgenic plants were crossed three times (construct *198*) or four times (constructs *195*, *196*, *162* and *175*) to the recurrent B73 parent line before phenotyping. During the introgression generations, plants were treated with a 2.5% Liberty solution applied to a single leaf to assay for 35SBAR gene-mediated resistance to Liberty herbicide (source, BASF). We also used transgene-specific genotype analyses to track integration events and determine transgene locus number by segregation analysis. DNA was made from leaf punches as previously described (Strable et al., 2017) and PCR-based genotype assays were performed using standard conditions with the primers described (**Table S3**). To genotype alleles at the endogenous *ZmRA1* locus in the presence of all but the *198* transgene, a CAPS assay was utilized to detect a SNP within with the *ra1-R* allele which results in the introduction of an *Acc*I restriction site. The 765 bp amplicon generated by primers RA8 and RA11 is digested by *Acc*I in *ra1-R* to generate two fragments, 334 bp and 431 bp. The *198* transgene contains the same *Acc*I SNP as *ra1-R*. Thus, in crosses with the *198* transgene, an additional *Msc*I dCAPS assay that detects the lesion in the *ra1-R* mutant allele was employed to distinguish the *198*-derived amplicons (i.e., without *Msc*I site to yield 190 bp) from the *ra1-R* derived amplicons (with the *Msc*I site to yield 155 bp and 35 bp following digestion).

Transgene events that segregated as single locus integrations and showed a stable herbicide-resistance phenotype were selected for qualitative or quantitative phenotyping analysis. To produce the segregating populations used for phenotyping *198*, *195* and *196*, plants heterozygous *ra1-R*/+ and hemizygous for the transgene of interest were crossed as females by *ra1-R*/*ra1-R* pollen of the recurrent B73 parent. To produce the *162* and *175* material for phenotyping we crossed females homozygous *ra1-R*/*ra1-R* and hemizygous for the transgene of interest by *ra1-R*/*ra1-R* pollen.

### Phenotypic analysis

All maize plant phenotyping was performed on field-grown plants in the summers of 2014 (constructs 195, 196 and 198) and 2018 (constructs 162 and 175), at the same location on the Woodruff Farm in Ames, Iowa. Tassel phenotype characters are summarized in **Fig. S5** and described here. Long branch zone was measured from the basal-most to the apical-most long branches. Central spike length was taken from the apical-most long branch to the tip of the tassel and comprised spikelet pairs. A long branch was defined as the typical basal long branches in maize, i.e. bearing only spikelet pairs, or as bearing a mix of spikelet pairs and single spikelets. Spikelet multimers were any branches bearing three or more single spikelets. Spikelet pair density was taken from a 1 cm band in circumference at the central spike midpoint.

### RNA *in situ* hybridization and expression analysis

Field-grown *S. bicolor* and growth chamber-grown *S. viridis* panicles were fixed overnight at 4° C in FAA. Samples were dehydrated through a graded ethanol series (50%, 70, 85, 95, 100) each one hour, with three changes in 100% ethanol. Samples were then passed through a graded Histo-Clear (National Diagnostics) series (3:1, 1:1, 1:3 ethanol: Histo-Clear) with 3 changes in 100% Histo-Clear; all changes were one hour each at room temperature. Samples were then embedded in Paraplast®Plus (McCormick Scientific), sectioned, and hybridized as described previously (Strable and Vollbrecht, 2019). Hybridizations were performed using antisense digoxygenin-labeled RNA probes to *ZmRA1* (**Table S3**) and *ZmKN1* (Jackson et al., 1994).

Field-grown, developmentally staged maize tassels were dissected away from leaf primordia and placed individually in 100 µL Trizol (Thermo-Fisher) and stored at -80°C in a 1.5 mL Eppendorf tube until processing. To process, 400 µL Trizol was added and tassel tissue was thawed and ground in the presence of Trizol using a plastic drill mount pestle. Total RNA was extracted as per the Trizol manufacturer and treated with RQ1 DNase (Promega) following the protocol outlined by the manufacturer, and converted to cDNA using RNA to cDNA EcoDry™ Premix (Double Primed) reagents (Takara Bio USA). The cDNA was diluted 1:1 with water, and 1.0 µL was used for PCR. PCR followed standard conditions using GoTaq®Green Master Mix (Promega corp.), Ta= 58°C, 1 min. extension at 72°C for 33 cycles. Primers are listed in **Table S3**.

### Conservation analysis of promoter *cis* sequences

For mVISTA analysis, genomic sequences (0.5 kb) upstream of the predicted 5’UTR regions of *RA1* in *Zea mays*, *Sorghum bicolor* and *Setaria viridis* were downloaded from https://ensembl.gramene.org and aligned using mVISTA LAGAN alignment (https://genome.lbl.gov/vista/mvista/submit.shtml). The plots depict 100 bp alignment windows at a similarity threshold 70% shaded in red.

To identify conserved non-coding sequences and binding motifs, the coding sequence of *Zea mays RA1* (Zm00001eb312340 – B73-REFERENCE-NAM-5.0) was used to find likely orthologs in other Panicoideae grasses. Sequences from *Chasmanthium laxum* (Chala.06G030500 – v1.1), *Miscanthus sinensis* (Misin03G169300 & MisinT268200 – v7.1), *Panicum halli* (Pahal.2G260300 – v3.2), *Paspalum vaginatum* (Pavag06G030400 – v.3.1), *Setaria viridis* (Sevir.2G209800 – v2.1) and *Sorghum bicolor* (Sobic.002G197700 and Sobic.002G197800 – v3.1.1) were identified using the BLAST tool in Phytozome v13. Sequence from *Coix lacryma-jobi* (Adlay0592-017T1) was selected from its own genome site. (http://phyzen.iptime.org/adlay/index.php). From all accessions, we took 2 kb upstream of the translation initiation site. First, conserved non-coding sequences from *RA1* promoter region sequence from Andropogoneae was determined using MEME (Bailey and Elkan, 1994). Then, the resulting motifs were searched in the other Panicoideae non-Andropogoneae grasses using FIMO (Grant et al., 2011). All sequences were compared against the non-redundant JASPAR CORE (2018) database from plants while using SEA to observe any possible well-known binding sites present internally (Bailey and Grant, 2021). The position of motifs from JASPAR were compared with the position of conserved non-coding sequences to check for overlap. Motifs from Clade ‘A’ ARFs were searched on the different sequences by using FIMO (Galli et al, 2018). Finally, these binding sites from the SEA analysis were used to search again in the Andropogoneae grasses using FIMO to obtain the relative coordinates in the *ZmRA1* promoter region (Grant et al., 2011).

### Accession numbers

*ZmRA1*, Zm00001eb312340; *ZmKN1*, Zm00001eb055920; *SbRA1^DS^*, Sobic.002G197700; *SbRA1^US^*, Sobic.002G197800; *SvRA1*, Sevir.2G209800; *ClRA1*, Chala.06G030500; *MsRA1*, Misin03G169300 & MisinT268200; *PhRA1*, Pahal.2G260300; *PvRA1*, Pavag06G030400; *Cl-j*, Adlay0592-017T1, *Eleusine coracana RA1* ELECO.r07.6AG0534810.1

## ACKNOWLEDGEMENTS

We are grateful to Brandi Sigmon for insightful discussion on *RA1* in sorghum and for comments on the manuscript. Additionally, we thank Pete Lelonek for assisting with greenhouse management and plant care. Many thanks to former undergraduate students, especially Emery Peyton, Charlie Beeler, Tryggve Rogers, Raven Saunders-Duckett, Matt Hirsch, Nicole Essner and Jack Schwickerath for their help with summer genetics nurseries and phenotypic analysis. We appreciate the insightful comments on the manuscript from Jack Satterlee. This work was supported by the National Science Foundation (IOS number 1238202 to E.V.). J.S. and A.A.R. are supported by North Carolina State University startup funds and USDA Hatch project 1026392.

## AUTHOR CONTRIBUTIONS

J.S., E.U.-W. and E.V. designed research; J.S., E.U.-W., S.B., and E.V. performed experiments; J.S., E.U.-W., A.A.R. and E.V. analyzed data; J.S., E.U.-W. and E.V. wrote the paper.

## Notes

### Competing Interest Statement

The authors have declared no competing interest.

