## Supplemental Tables ST1 - ST3 for "Interspecies transfer of syntenic *RAMOSA1* orthologs and promoter *cis* sequences impacts maize inflorescence architecture"

**Supplemental Table 1. Reference genome coordinates, and construct and trangenic event information**

| Description | Construct | RefGen Coordinates[1] | size (bp) | Upstream (bp)[2] | CDS (bp) | Downstream (bp)[3] | Events[4] | Complementation[5] |
|---|---|---|---|---|---|---|---|---|
| *ZmRA1* | 198 | *Zm Chr7:114955807-114961642* | 5834 | 2958 | 528 | 2348 | 5 | Y, Y, Y, Y, Y-p |
| *SbRA1 TAN* | 195 | *Sb Ch2: 58694932-58700945* | 6026 | 722<br>2027* | 548 US<br>555 DS | 2027*<br>2174 | 3 | Y-p, Y-p, Y-p, Y-p |
| *SbRA1 DS* | 196 | *Sb Ch2: 58694932-58699332* | 4404 | 1675 | 555 | 2174 | 3 | Y, Y, Y-p |
| *SvRA1* | 162 | *Sv Ch2: 29092485-29096480* | 4028 | 1537 | 522 | 1969 | 4 | Wk-p, N, N, N |
| *pZmRA1::SvRA1* | 175 | *Zm Chr7:114955807-114958766*<br>*Sv Ch2: 29092485- 29094453* | 2958<br>2491 | 2958<br>np | np<br>522 | np<br>1969 | 1 | Y |

np, not present

[1] Zm=Zea mays B73 v5,Sb=Sorghum bicolor NCBIv3, Sv=Setaria viridis v2.0

[2] Sequence upstream (5-prime) of the CDS start codon, therefore includes 5-prime UTR

[3] Sequence starts immediately after CDS stop codon position, therefore includes 3-prime UTR.

[4] Number of independent transgenic events with a stably heritable, apparently single-locus, BAR-expressing transgene.

[5] Summary of phenotyping, degree of correction of ra1-R in transgenics; one comma-separated entry for each event; see text and Table S2 for details.

[5] Summary of phenotyping, continued: Y, Yes; Wk, weak or partial; N, No; p, pleiotropy.

* DNA between position of non-frameshifted stop codon position in SbUS and the start codon of SbDS

**Supplemental Table 2. Transgene genetic segregation data**

| Event[1] | Season[2] | Generation[3] | TG+ | NTG | Total | p-value[4] | Qualitative phenotype[5] |
|---|---|---|---|---|---|---|---|
| 198.7.3* | Su11 | BC3 | 68 | 65 | 133 | 0.795 | Y |
| 198.23.2 | Su11 | BC3 | 20 | 19 | 39 | 0.873 | Y |
| 198.23.4 | Su11 | BC3 | 68 | 85 | 153 | 0.169 | Y |
| 198.27.3 | Su11 | BC3 | 149 | 160 | 309 | 0.531 | Y-p |
| 198.32.3 | Su11 | BC3 | 40 | 38 | 78 | 0.821 | Y |
| 195.8.3* | Su14 | BC4 | 117 | 124 | 241 | 0.652 | Y-p |
| 195.20.4* | Su11, GH14 | BC3 | 35 | 30 | 65 | 0.535 | Y-p |
| 195.46.1* | Su11, GH14 | BC3 | 39 | 30 | 69 | 0.279 | Y-p |
| 195.14.4 | Su11 | BC3 | 72 | 74 | 146 | 0.869 | Y-p |
| 196.19.2* | Su11, GH14 | BC3 | 33 | 35 | 68 | 0.808 | Y |
| 196.7.3* | Su11, GH14 | BC3 | 38 | 33 | 71 | 0.553 | Y |
| 196.14.5* | Su14 | BC4 | 120 | 113 | 233 | 0.647 | Y-P |
| 162.26.1* | Su17, Su18 | BC4 | 82 | 98 | 133 | 0.233 | Wk-p |
| 162.10.6.5 | Su17 | BC3 | 22 | 19 | 41 | 0.639 | N |
| 162.28.2.7 | Su18 | BC4 | 58 | 51 | 110 | 0.503 | N |
| 162.7.3 | Su17 | BC4 | 19 | 13 | 32 | 0.289 | N |
| 175.7* | Su18 | BC3 | 141 | 128 | 220 | 0.428 | Y |

TG+, plants containing transgene based on bialophis resistence phenotype

NTG, non-transgenic plants based on bialophis resistence phenotype

[1] Transgene events phenotyped in detail are marked by *

[2] Su, Summer; GH, greenhouse; two-digit numerals xy refer to year 20xy

[3] BCn, backcrossed to B73 n generations

[4] p-value, chi-square test, 1:1 segregation

[5] Qualitative phenotype score, degree of correction of *ra1-R* in TG+ individuals; see text for additional details.

[5] Qualitative phenotype score, continued: Y, Yes; Wk, weak or partial; N, No; p, pleiotropy.

**Supplemental Table 3 . Primers used in this study**

| Name | Sequence | Description | Purpose |
|---|---|---|---|
| RA72 | CATTAATGAAATTTATTCGTCGTCAAGC | *ZmRA1* locus | Vector : 198 |
| RA76 | AAGCTTACACAGTAACACGGGTGC | *ZmRA1* locus | Vector: 198 |
| EUO237 | CACACGCGCGCCACTCGACTAG | SNP introduction in *ZmRA1* | Vector: 198 |
| EUO238 | GTGTGAAGTATACTGTTGGTGGATCAGTCTG | SNP introduction in *ZmRA1* | Vector: 198 |
| EUO239 | CAGTATACTTCACACCGTATTGCTGCTCC | SNP introduction in *ZmRA1* | Vector: 198 |
| EUO240 | CTTCCCCTCGACGTCCTTGGTCG | SNP introduction in *ZmRA1* | Vector: 198 |
| EUO231 | CTGGACTTCAGCCTGCCGGTACCGCC | *Hin* dIII site 3' BAR | Vector: pSB11_BAR |
| EUO232 | CAAGCTTGTAAAACGACGGCCAGTGCCAAG | *Hin* dIII site 3' BAR | Vector: pSB11_BAR |
| EUO233 | GAAGCTTGCACACATCCAACAACGACACAG | *Hin* dIII site for *SbRA1* DS | Vector: 195 |
| EUO234 | GCTACAGGACCTGCACCCATGCATG | *Hin* dIII site for *SbRA1* DS | Vector: 195 |
| EUO421 | CGGCTTGCTGGTCTAAGTCTAACTACCTTG | *SvRA1* locus | Vector: 162 |
| EUO423 | CATGCAAGCTGGGGATCCGCAGCCTGTGGCTCGCCTGTCCG | *SvRA1* locus | Vector: 162 |
| EUO424 | TAGACCAGCAAGCCGCCTATATGGAGAGAGATG | *SvRA1* locus | Vector: 162 |
| EUO426 | GCCACTCAGCAAGCTTCTGCGCGCGCTTTCGCTTTG | *SvRA1* locus | Vector: 162 |
| EUO482 | CATGCAAGCTGGGGATCCGCAG | *SvRA1* CDS and 3' flank | Vector: 175 |
| EUO489 | GACTAGCTAGCAGCTATGGAGAGAGATGATGGCTACACCA | *SvRA1* CDS and 3' flank | Vector: 175 |
| EUO479n | AGGAGCCACTCAGCAAGCTTACACAGTAACAC | *ZmRA1* 5' promoter region | Vector: 175 |
| EUO488 | ATCATCTCTCTCCATAGCTGCTAGCTAGTCGAGTGGCG | *ZmRA1* 5' promoter region | Vector: 175 |
| EUO318 | AGCACCATGCGCTAGCAGCGAC | *ra1-R* | *Msc* I dCAPS assay |
| EUO317 | AAGAAGGAGTTCAGATCAGCACAAGGGCTGGGTGG | *ra1-R* | *Msc* I dCAPS assay |
| RA8 | TGCTCTATCTTGCCTCTTCATGC | *ZmRA1* locus | *Acc* I CAPS assay |
| RA11 | TGCACTGCACGTACCCATTGTAG | *ZmRA1* locus | *Acc* I CAPS assay |
| SbRa1-F2 | TCCACCAGCAGTACATGTCG | *SbRA1* DS | 195,196; RT-PCR |
| EUO386x | CAAGTTCTTCACCTCGGAGTAGTAG | *SbRA1* DS | 195,196 ;RT-PCR |
| EUO411 | CAGCAAGCCGCCTATATGGAG | *SvRA1* locus | genotype 162 |
| EUO412 | TCGTCTCAGGAGTGGCCAAGTC | *SvRA1* locus | genotype 162 |
| ZmRA1-F2 | CTGCCGCCACAGGTAAGG | *ZmRA1* | RT-PCR |
| ZmRA1-R2 | CCCTCGACGTCCTTGGTC | *ZmRA1* | RT-PCR |
| ZmRA1-F1 | GCCGCCACAGGTAAGGTC | *ZmRA1 in situ* | probe amplicon |
| ZmRA1-R3 | TTCCCCTCGACGTCCTTG | *ZmRA1 in situ* | probe amplicon |
| ZmUbi-R1 | CTGAAAGACAGAACATAATGAGCACAGGC | *ZmUbiquitin1* | *RT-PCR* |
| ZmUbi-F1 | TAAGCTGCCGATGTGCCTGCGTCG | *ZmUbiquitin1* | *RT-PCR* |
