## Supplemental figures S1 - S10 for "Interspecies transfer of syntenic *RAMOSA1* orthologs and promoter *cis* sequences impacts maize inflorescence architecture"

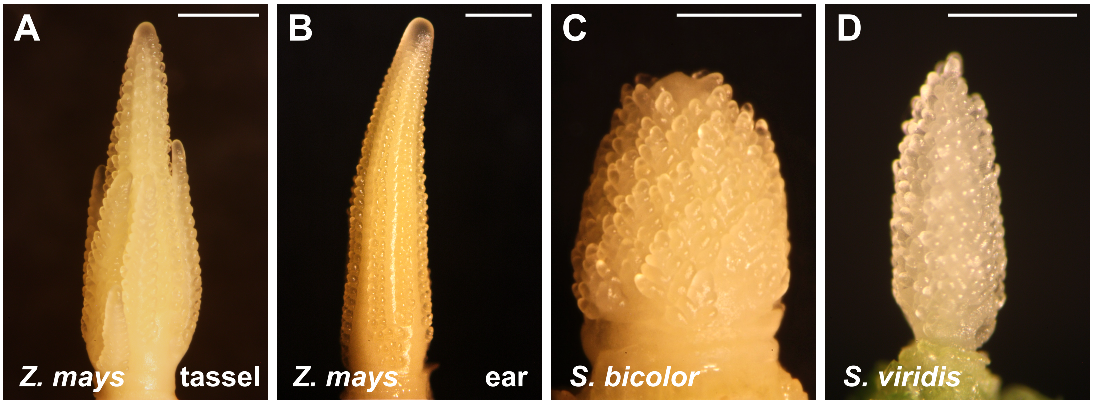


**Figure S1. Developing inflorescences of maize, sorghum and *S. viridis*.** Dissected inflorescences representing (A) maize tassel, (B) maize ear, (C) sorghum panicle, and (D) *S. viridis* panicle. Scale bars, 1 mm (A-C); 0.5 mm (D).

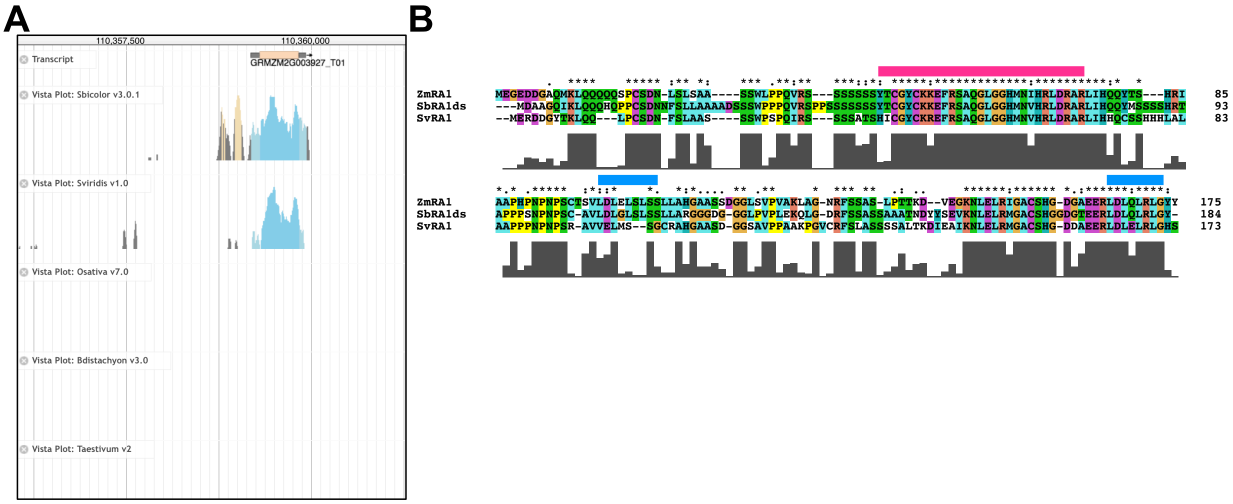


**Figure S2. *RA1* locus and amino acid sequence in Panicoid grass species.** (A) mVISTA plots of *RA1* syntenic genomic regions in *Zea mays, Sorghum bicolor, Setaria viridis, Oryza sativa, Brachypodium distachayon* and *Triticum aestivum* showing that the *RA1* locus is present only in the *Zea mays, Sorghum bicolor,* and *Setaria viridis* genomes. Coding and non-coding regions that share >70% identity are shown as blue and tan peaks, respectively. (B) Amino acid alignment of RA1 from *Z. mays*, *S. bicolor* downstream (DS) copy and *S. viridis* rendered by ClustalX. Horizonal magenta line, $C_2H_2$ zinc finger domain. Horizonal blue line, EAR motif.

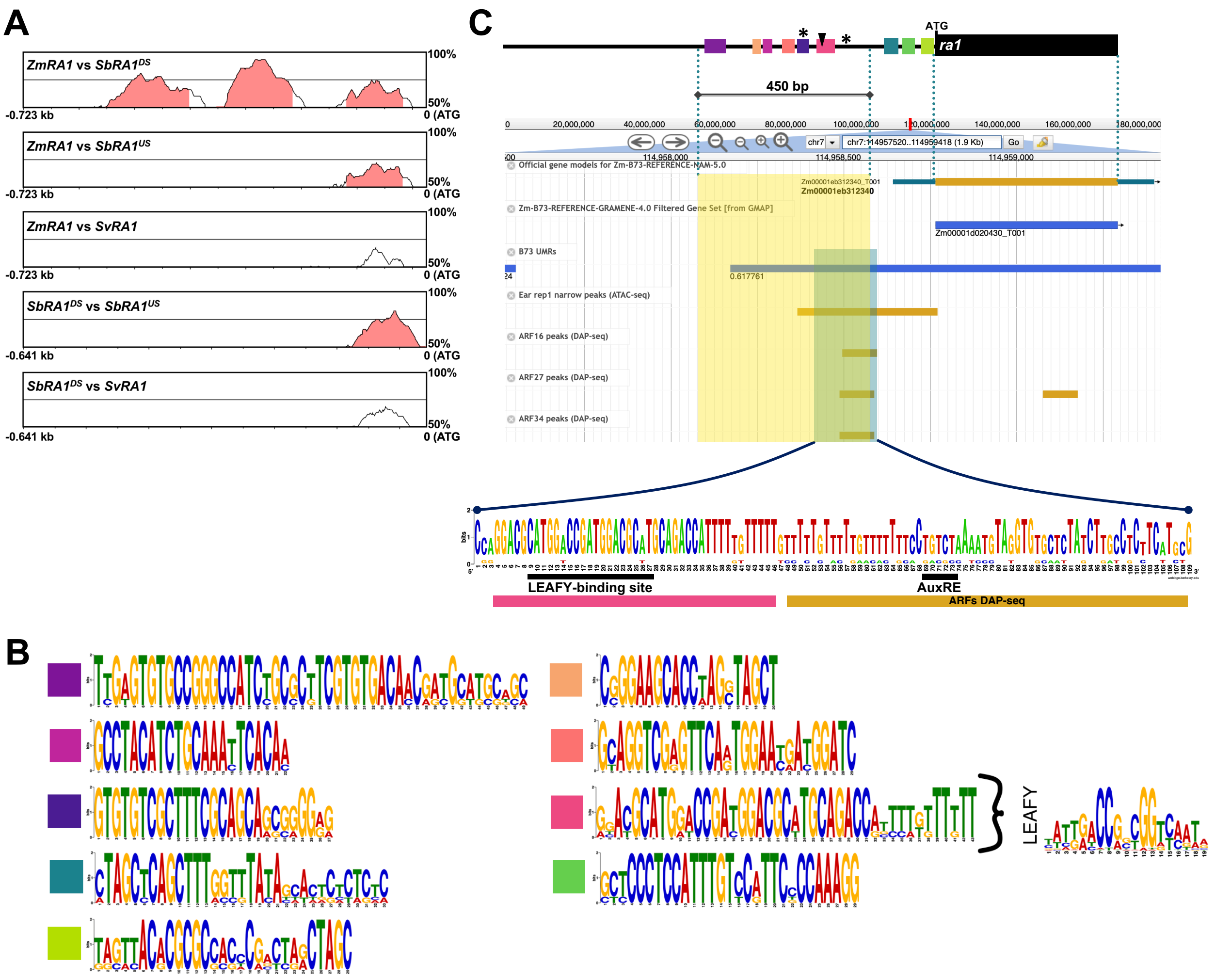


**Figure S3. Conserved *cis* sequences in *RA1* promoter regions of Panicoid grasses.** (A) mVISTA plots comparing *RA1* near-promoter regions of maize, sorghum upstream, sorghum downstream and setaria show conserved regions among maize *RA1* and the sorghum copies. Red regions of mVISTA plots indicate >70% sequence similarity over 100-bp windows. Position 0 is the first base upstream of the ATG codon. (B) Sequence logos of the conserved motifs (see Figures 1H and S3C and Supplemental Dataset) present in the *RA1* promoter region of the Andropogoneae grasses. The LFY motif is indicated next to the conserved motif it resides in. (C) Cartoon depiction of the *ZmRA1* promoter region with conserved non-coding *cis* sequences, and its coding region. To scale below is a genome browser view of the Zm-B73-Reference-NAM-5.0 genome assembly displaying the following tracks (from top to bottom): *ZmRA1* locus; B73 unmethylated regions (UMRs); Ear rep 1 narrow peaks (ATAC-seq); Leaf rep 1 narrow peaks (ATAC-seq); ZmARF16 peaks (DAP-seq); ZmARF27 peaks (DAP-seq); ZmARF34 peaks

(DAP-seq). Area highlighted in yellow corresponds to the block of six conserved non-coding *cis* sequences in the distal *ZmRA1* promoter region. Area highlighted in blue corresponds to the one conserved non-coding *cis* sequence block and ARF DAP-seq peaks that house LFY and AuxRE binding motifs, respectively. Sequence logo in (C) derived from *ZmRA1* and *SbRA1*$^{DS}$.

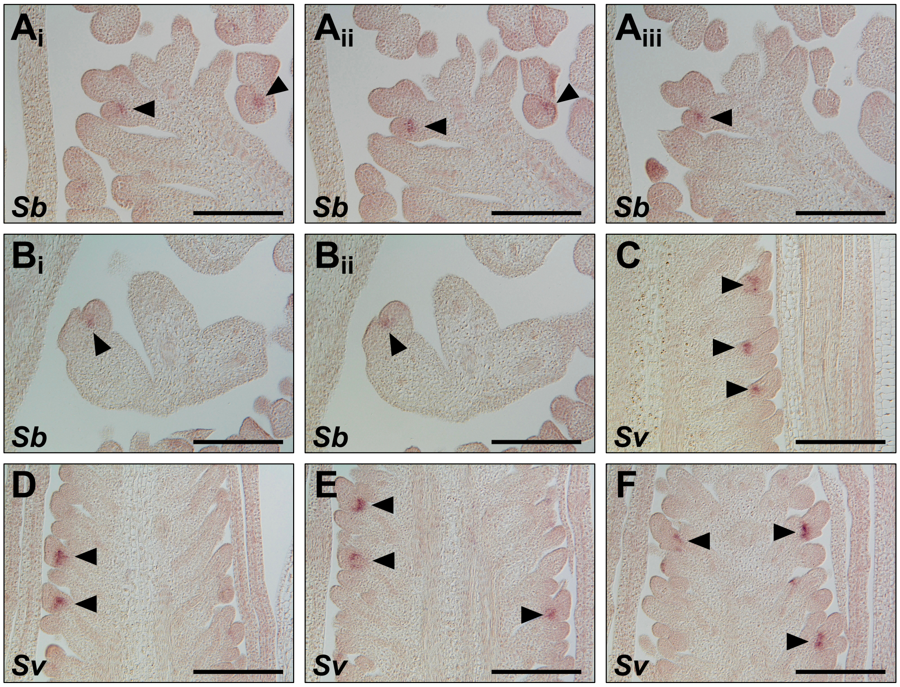


**Figure S4. RNA *in situ* hybridization in sorghum and *S. viridis* inflorescences.** Antisense RNA probes to *ZmRA1* (A-F) were hybridized to longitudinal sections of developing inflorescences from sorghum, *Sb* (A, B) or *S. viridis, Sv* (C-F). $A_i$-$A_{iii}$ and $B_i$-$B_{ii}$ represent serial sections. Arrowheads denote *RA1* transcript accumulation. Scale bars, 100 µm.

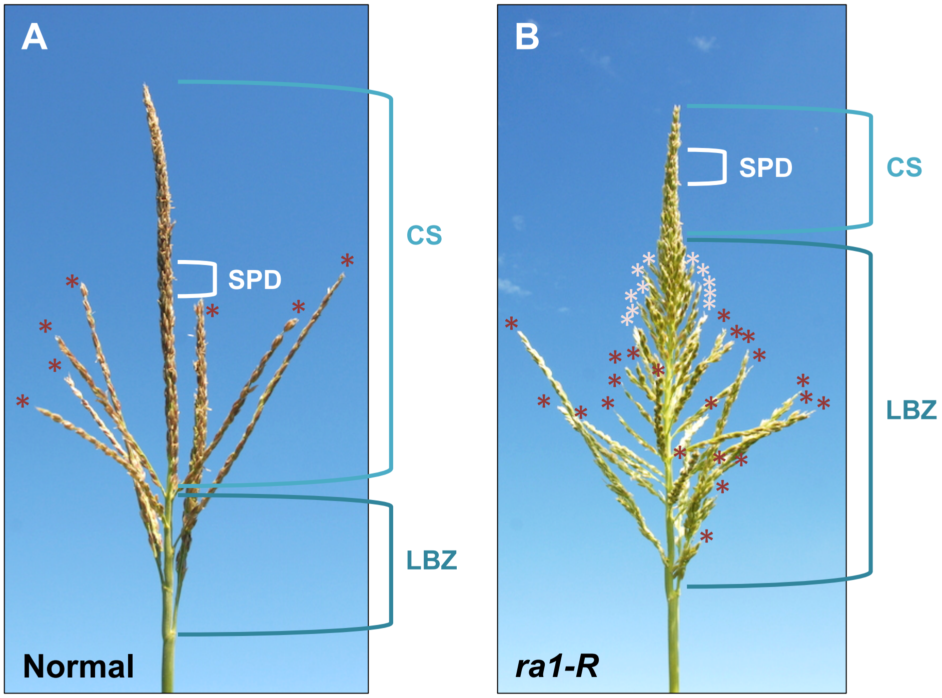


**Figure S5. Tassel traits quantified in this study.** Field-grown normal (A) and *ra1-R* (B) tassels. Long branch zone was measured from the basal-most to the apical-most long branches. Central spike length was taken from the apical-most long branch to the tip of the tassel and comprised spikelet pairs. A long branch was defined as the typical basal long branch in maize, i.e. bearing only spikelet pairs, or as bearing a mix of spikelet pairs and single spikelets. Spikelet multimers were any branches bearing three or more single spikelets. Spikelet pair density taken from a 1 cm band in circumference at the central spike midpoint. LBZ, long branch zone; CS, central spike; SPD, spikelet pair density. Dark red asterisk, Long branches (LBs); light red asterisks, spikelet multimers.

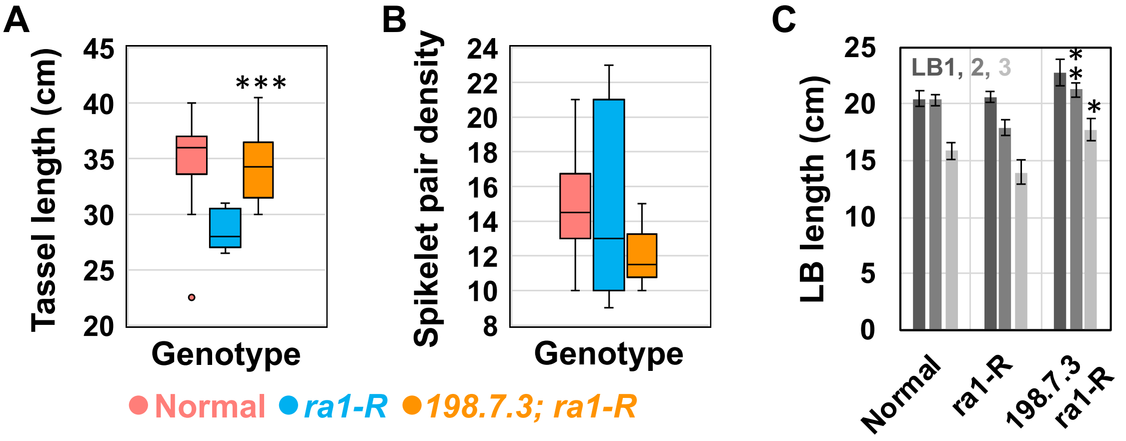


**Figure S6. Tassel traits in *ra1-R* mutants expressing the *198* transgene cassette**. (A) Tassel length (B) Spikelet pair density from a 1 cm band in circumference at the CS midpoint. (C) Long branch (LB) length for the three basal-most LBs. (A, B) Box and whisker plots, the bottom and top boxes represent the first and third quartile, respectively, the middle line is the median, and the whiskers represent the minimum and maximum values, outlier data points are displayed as individual dots. (C) Mean ± SD; Two-tailed Student's *t* test for transgene vs. *ra1-R* ***$P$<0.001, **$P$<0.01, *$P$<0.05; normal, n = 20; *ra1-R*, n = 10; 198.7.3, n = 8.

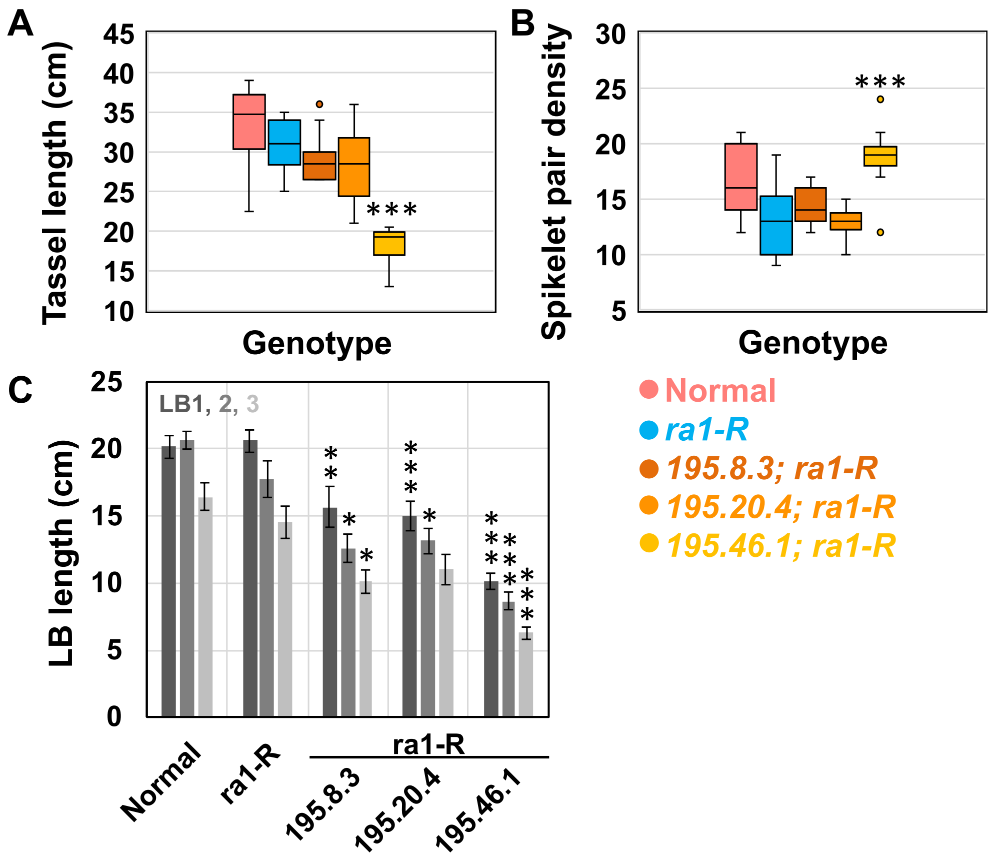


**Figure S7. Tassel traits in *ra1-R* mutants expressing the *195* transgene cassette**. (A) Tassel length (B) Spikelet pair density from a 1 cm band in circumference at the CS midpoint. (C) Long branch (LB) length for the three basal-most LBs. (A, B) Box and whisker plots, the bottom and top boxes represent the first and third quartile, respectively, the middle line is the median, and the whiskers represent the minimum and maximum values, outlier data points are displayed as individual dots. (C) Mean ± SD; Two-tailed Student's *t* test for transgene vs. *ra1-R* ***$P<0.001$, **$P<0.01$, *$P<0.05$; *ra1-R*, n = 18; 195.8.3, n = 11; 195.20.4, n = 12; 195.46.1, n = 11.

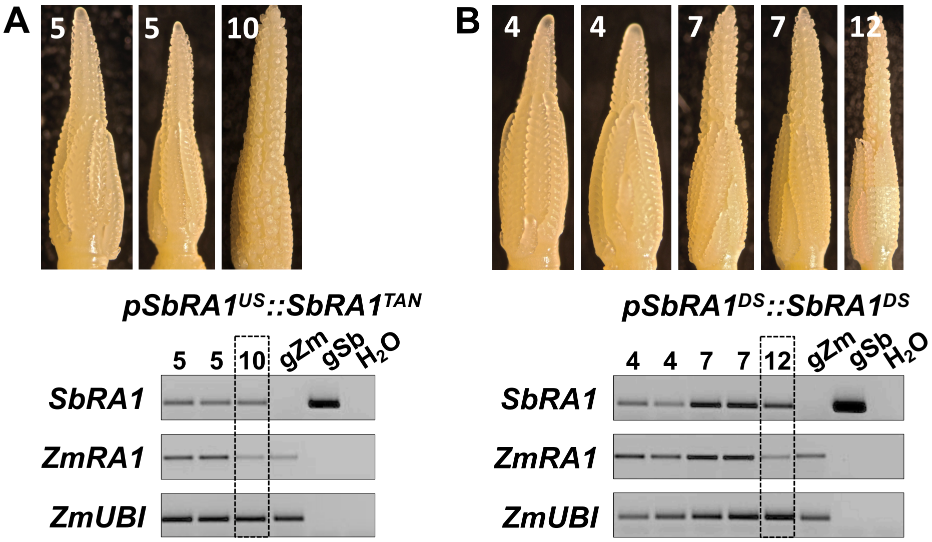


**Figure S8. Expression of *195* and *196* transgenes and endogenous *ZmRA1* in developing tassels.** Reverse transcription followed by semi-quantitative PCR of individual tassels expressing *195.20.4* (A) or *196.14.5* (B) transgenes, and endogenous *ZmRA1* and *ubiquitin* (*UBI*) at the developmental stages (tassel lengths in mm) labeled within each tassel image. Data are for 33 PCR cycles; genomic DNA positive control, water negative control.

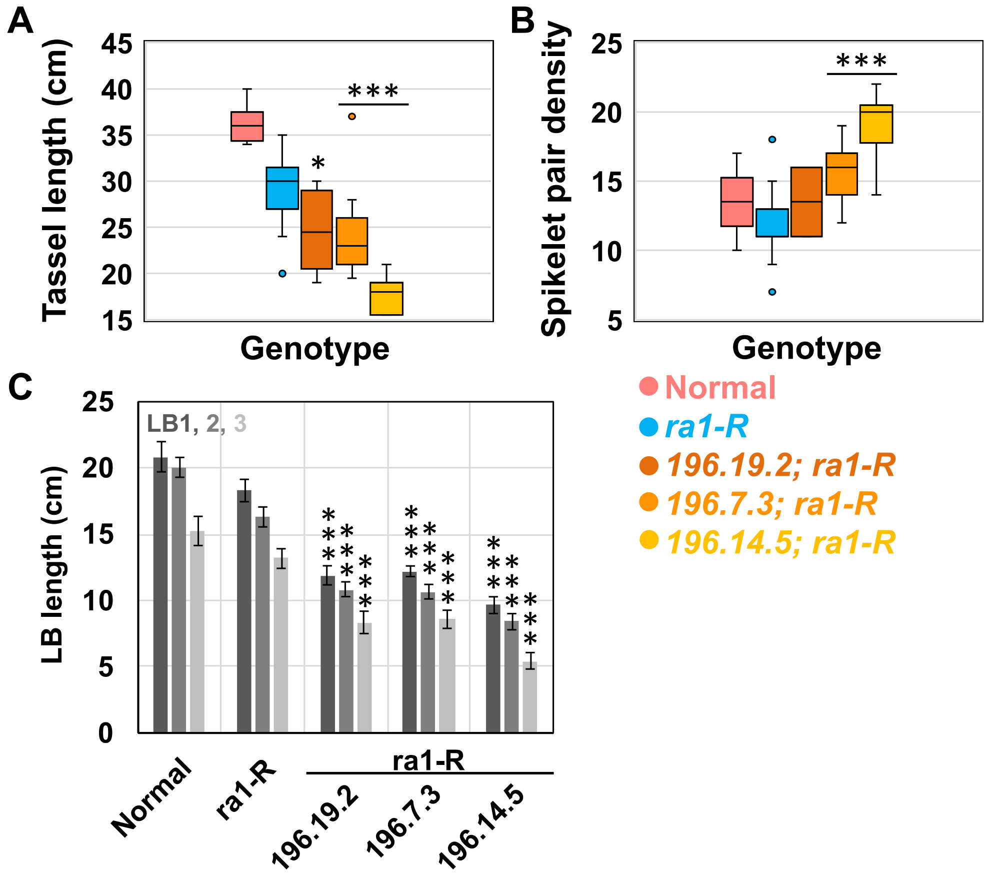


**Figure S9. Tassel traits in *ra1-R* mutants expressing the *196* transgene cassette**. (A) Tassel length (B) Spikelet pair density from a 1 cm band in circumference at the CS midpoint. (C) Long branch (LB) length for the three basal-most LBs. (A, B) Box and whisker plots, the bottom and top boxes represent the first and third quartile, respectively, the middle line is the median, and the whiskers represent the minimum and maximum values, outlier data points are displayed as individual dots. (C) Mean ± SD; Two-tailed Student's *t* test for transgene vs. *ra1-R* ****P*<0.001, **P*<0.05; *ra1-R*, n = 15; 196.19.2, n = 10; 196.7.3, n = 12; 196.14.5, n = 10.

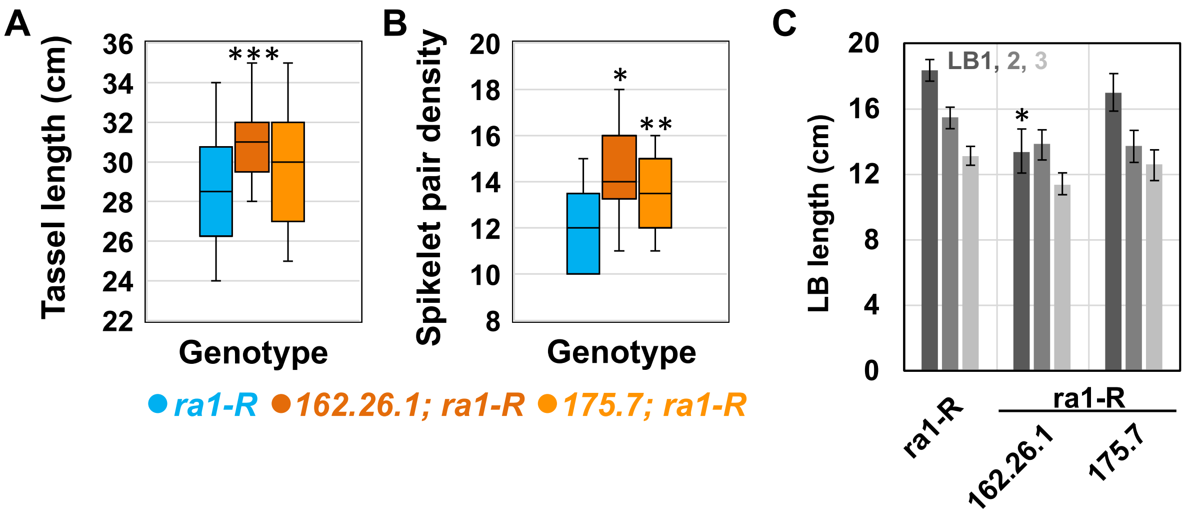


**Figure S10. Tassel traits in *ra1-R* mutants expressing either the *162* or *175* transgene cassette**. (A) Tassel length (B) Spikelet pair density from a 1 cm band in circumference at the CS midpoint. (C) Long branch (LB) length for the three basal-most LBs. (A, B) Box and whisker plots, the bottom and top boxes represent the first and third quartile, respectively, the middle line is the median, and the whiskers represent the minimum and maximum values, outlier data points are displayed as individual dots. (C) Mean ± SD; Two-tailed Student's *t* test for transgene vs. *ra1-R* ***$P$<0.001, **$P$<0.01, *$P$<0.05; *ra1-R*, n = 15; 162.26.1, n = 12; 175.7, n = 9.
